# Prolonging lung cancer response to EGFR inhibition by targeting the selective advantage of resistant cells

**DOI:** 10.1101/2023.06.19.545595

**Authors:** Lisa Brunet, David Alexandre, Jiyoung Lee, Maria del Mar Blanquer-Rosselló, Alexis Guernet, Houssein Chhouri, Zoulika Kherrouche, Arnaud Arabo, Shen Yao, David Godefroy, Julie Dehedin, Jian-Rong Li, Céline Duparc, Philippe Jamme, Audrey Vinchent, Caroline Bérard, David Tulasne, Sabrina Arena, Alberto Bardelli, Chao Cheng, Byoung Chul Cho, Cédric Coulouarn, Stuart A. Aaronson, Alexis B. Cortot, Youssef Anouar, Luca Grumolato

## Abstract

Non-small cell lung cancers (NSCLCs) treated with tyrosine kinase inhibitors (TKIs) of the epidermal growth factor receptor (EGFR) almost invariably relapse in the long term, due to the emergence of subpopulations of resistant cells. Here we show that the lack of sensitivity of these cells to EGFR-TKIs constitutes a vulnerability that can be potentially targeted. Through a DNA barcoding approach, we demonstrate that the clinically approved drug sorafenib specifically abolishes the selective advantage of EGFR-TKI-resistant cells, while preserving the response of EGFR-TKI-sensitive cells, thus resulting in overall inhibition of clonal evolution within the tumor cell mass population. Sorafenib is active against multiple mechanisms of resistance/tolerance to EGFR-TKIs and its effects depend on early inhibition of MAPK interacting kinase (MNK) activity and signal transducer and activator of transcription 3 (STAT3) phosphorylation, and later down-regulation of MCL1 and EGFR. Using several xenograft and allograft models to recapitulate different mechanisms and kinetics of acquired resistance, we show that the sorafenib-EGFR-TKI combination can substantially delay tumor growth and promote the recruitment of inflammatory cells. Together, our findings indicate that sorafenib can substantially prolong the response to EGFR-TKIs by targeting NSCLC capacity to adapt to treatment through the emergence of resistant cells.

## INTRODUCTION

Clonal driver mutations are a target of choice for cancer therapy. One of the first and more successful examples of this type of treatment is represented by EGFR-TKIs, including first generation gefitinib and erlotinib, and third generation osimertinib, which are used as first line treatment for advanced EGFR-mutant NSCLCs (Peters et al., 2021). Despite objective response rates that can exceed 80% (Ramalingam et al., 2018; Soria et al., 2018b), EGFR-TKIs almost invariably fail in the long term because of the development of resistance (Passaro et al., 2021). Current guidelines are based on a sequential scheme, in which EGFR-TKIs are administered as single agents until progression, followed by treatment with chemotherapy or, when an actionable mechanism of resistance is identified, combinations with another targeted drug (Sequist et al., 2020; Hanna et al., 2021).

In recent years, a better understanding of how cancer can adapt to treatment and become insensitive has suggested that, instead of waiting for tumor relapse, therapies could be designed to anticipate and delay the onset of resistance, thus prolonging EGFR-TKI response. Tumors are formed by an intricate combination of heterogeneous subclonal populations that evolve in the presence of a selective pressure, such as therapeutic intervention (Vasan et al., 2019). As for other targeted drugs, EGFR-TKI treatment provokes over time the emergence of resistant clones, which can be present before the therapy begins or originate *de novo* from pools of persister/tolerant cells (Bhang et al., 2015; Hata et al., 2016; Ramirez et al., 2016; Shen et al., 2020), eventually resulting in tumor relapse. Different drug combinations have been proposed to interfere with these processes and inhibit the acquisition of resistance (Jin et al., 2023). Some of them are meant to enhance inhibition of EGFR signaling by targeting other components of the pathway, using compounds that act synergistically with EGFR-TKIs (Tricker et al., 2015). Other strategies are designed to specifically block pathways involved in the acquisition of a tolerant/persister phenotype (Hata et al., 2016; Shah et al., 2019; Kurppa et al., 2020; Tanaka et al., 2021; Noronha et al., 2022). However, among the different combinations that have been clinically tested in treatment naïve patients (Passaro et al., 2021), so far the only one that has shown a significant increase in overall survival (OS) is with chemotherapy. Indeed, while previous studies in unselected patients failed to demonstrate such a benefit, two recent independent phase III trials in EGFR-mutant NSCLC patients established a clear superiority for the association of gefitinib with carboplatin and pemetrexed, compared to gefitinib alone, albeit at the expense of a higher toxicity (Hosomi et al., 2020; Noronha et al., 2020). By definition, chemotherapy targets proliferating cells, so it is conceivable that, while the majority of cancer cells are inhibited by gefitinib, carboplatin and pemetrexed exert a stronger effect on the subpopulations cells whose growth is not, or only poorly affected by the EGFR-TKI. This combination could then function, at least in part, by preventing the emergence of gefitinib resistant cells. These observations suggest that: i) tumor capacity to evolve and adapt to therapy can be inhibited by neutralizing the selective advantage of resistant over sensitive cells; ii) the capacity of resistant cells to grow in the presence of EGFR-TKIs could represent, by itself, some kind of collateral vulnerability that can be pharmacologically targeted; iii) since it’s not aimed at a specific mechanism of resistance, an approach directed against this type of vulnerability could be more broadly effective in prolonging the response to EGFR-TKIs.

Screens for new drug combinations are generally designed to find compounds that act synergistically. This type of approach is not suited to identify molecules capable of inhibiting the selective advantage of resistant cells in the presence of EGFR-TKIs. We previously devised a DNA barcoding strategy to generate and track small pools of NSCLC resistant cells within a mass population of sensitive cells. We showed that EGFR-TKIs induce a rapid enrichment of the barcoded cells, and this effect can be blocked through specific inhibition of the mechanism of resistance (Guernet et al., 2016). Through this approach, here we showed that, in combination with gefitinib, pemetrexed does not further inhibit the growth of EGFR-TKI sensitive cells, but it prevents the emergence of resistant cells. To identify other drugs capable of exerting a similar effect with potentially lower toxicity, we performed an unbiased functional screen using a NSCLC cell model containing three distinct mechanisms of resistance to EGFR-TKIs. We found that the clinically approved multikinase inhibitor sorafenib can prevent the emergence of cells displaying various types of resistance/tolerance to EGFR-TKIs, including secondary EGFR mutation, aberrant activation of different downstream components of EGFR signaling, amplification/overexpression of other receptor tyrosine kinase and epithelial to mesenchymal transition (EMT). Consistent with a specific vulnerability of EGFR-addicted cancer cells to sorafenib, gene expression analysis of tumor samples from various independent clinical trials revealed that patient response to this drug is significantly correlated with a high EGFR transcriptional score. Mechanistically, we demonstrated that the effects of sorafenib in NSCLC cells are independent of the MAPK pathway and rely instead of early inhibition of MNK activity and STAT3 phosphorylation, and later down-regulation of MCL1 and EGFR. Finally, using different xenograft and allograft models of acquired resistance to osimertinib, we showed that the osimertinib-sorafenib combination can promote the recruitment of inflammatory cells and substantially prolong the effects of EGFR-TKI treatment *in vivo*, even in extremely aggressive and rapidly progressing tumors.

## RESULTS

### Similarly to pemetrexed, sorafenib inhibits the selective advantage of resistant NSCLC cells in the presence of EGFR-TKIs

Co-treatment with chemotherapy was recently shown to prolong NSCLC response to gefitinib (Hosomi et al., 2020; Noronha et al., 2020). We speculated that in these patients gefitinib sensitive cancer cells are mostly inhibited by the EGFR-TKI, while chemotherapy acts on the subpopulations of resistant cells capable of growing in the presence of gefitinib. To test this hypothesis, we investigated the effects of pemetrexed and gefitinib, alone or in combination, on the viability of human NSCLC PC9 cells. While only pemetrexed inhibited the growth of gefitinib resistant cells (Figure S1A), the two drugs didn’t exert any additive/synergistic effects in parental cells (Figure 1A), supporting the notion that chemotherapy mainly affects the growth of cells that don’t respond to EGFR-TKIs. We devised a strategy, named CRISPR-barcoding, to generate and trace resistant clones within a population of EGFR-TKI sensitive cells (Guernet et al., 2016). As shown in Figure 1B, gefitinib treatment provoked an increase in the fraction of cells containing the resistance mutation EGFR-T790M, and this effect was blocked by co-treatment with pemetrexed. Together, these data strongly suggest that the clinical benefit of the EGFR-TKI/chemotherapy combination depends, at least in part, on its capacity to prevent or delay the emergence of EGFR-TKI-resistant cells. This association significantly prolonged the response of EGFR-mutant NSCLCs, but it also increased the rate of adverse effects (Hosomi et al., 2020; Noronha et al., 2020). We performed a small molecule screen to identify other drugs capable of inhibiting the emergence of cells containing multiple mechanisms of resistance to EGFR-TKI, while potentially displaying lower toxicity. We used a CRISPR-barcoding cell model in which three different small pools of cells bearing a distinct resistance mutation are intermingled within a mass population of sensitive cells. As expected, the third generation EGFR-TKI WZ42002 and the ALK-TKI TAE684 specifically blocked the enrichment of the EGFR-T790M and EML4-ALK subpopulations, respectively, but they had no effect on the other resistant cells. Intriguingly, we found that sorafenib could prevent the emergence of all three mechanisms of resistance at *in vitro* concentrations that were clinically relevant (Hu et al., 2008; Liston and Davis, 2017) (Table S1, Figure S1B-C). Sorafenib is a multikinase inhibitor targeting RAF, the vascular endothelial growth factor receptors (VEGFR), platelet-derived growth factor receptors (PDGFR), KIT and RET, and it has been approved for the treatment of advanced kidney, thyroid and liver cancers (Wilhelm et al., 2006). Of note, a phase III trial has previously shown that a sorafenib monotherapy can improve OS of patients with EGFR mutant, but not wild-type (wt) NSCLC (Paz-Ares et al., 2015). Consistent with these data, a transcriptional signature to detect EGFR activation was used to predict the sensitivity to sorafenib in both lung and liver cancer patients (Cheng et al., 2020). We found one other publicly available dataset containing gene expression and clinical data of cancer patients treated with sorafenib (Gudkov et al., 2022). As shown in Figure 1C, renal cell carcinomas that responded to this drug displayed higher EGFR-scores compared to non-responders. Together, these observations indicate that tumors with activation of EGFR signaling are sensitive to sorafenib, thus supporting the notion that this drug could be a promising candidate to prolong EGFR-TKI response.

**Figure 1.**
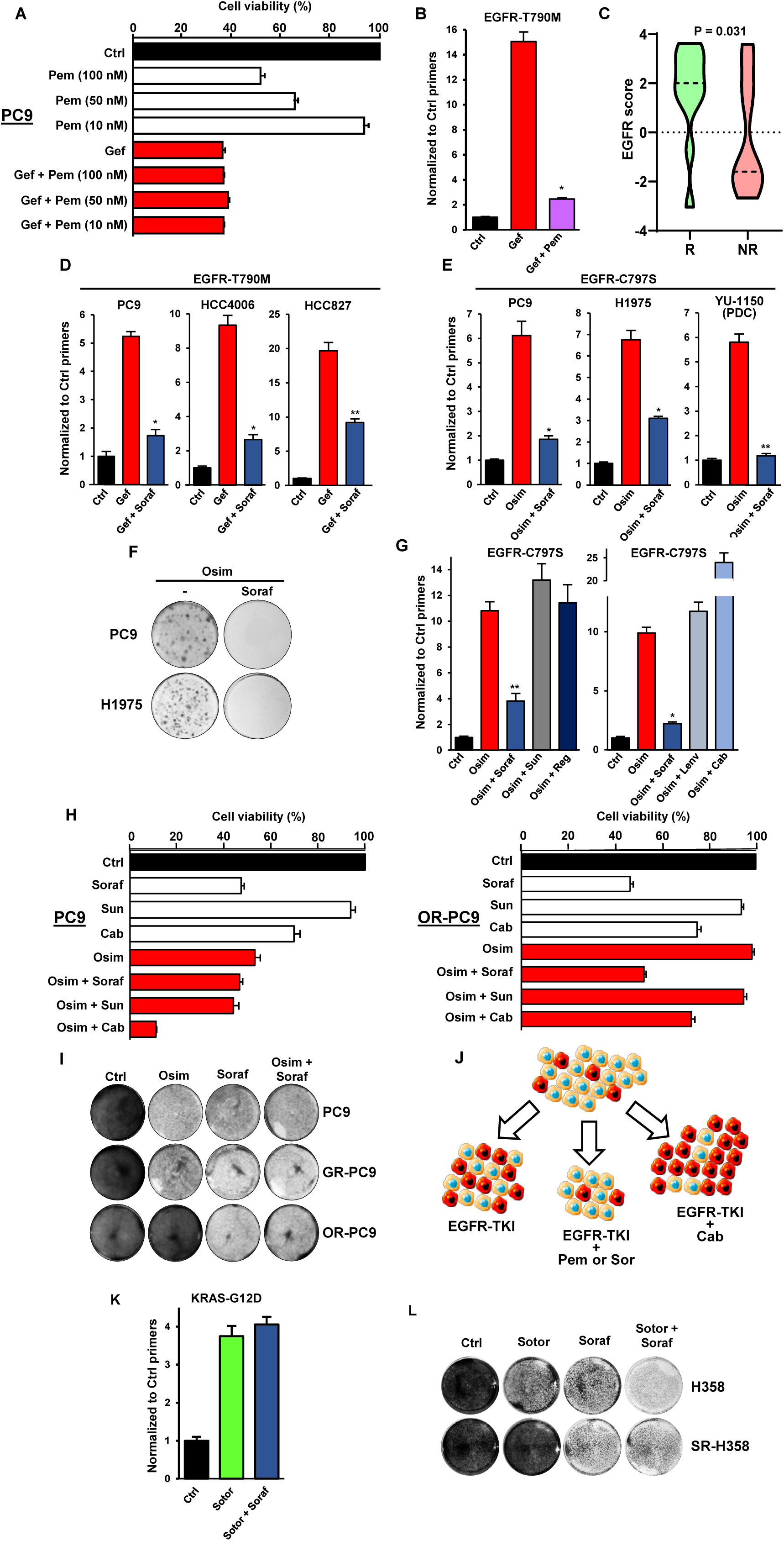
Pemetrexed and sorafenib inhibit the emergence of resistant subpopulations of NSCLC cells induced by EGFR-TKIs. **A.** Cell viability assay of PC9 cells treated for 5 days with pemetrexed (100, 50 or 10 nM; Pem) and gefitinib (1 µM; Gef), alone or in combination. The fraction of viable cells was measured by CellTiter-Glo and normalized to the DMSO treated control. The mean ± SEM (n=6) of one representative of three independent experiments is shown. **B.** A mass population of PC9 cells containing a small pool of cells bearing the EGFR-T790M mutation generated by CRISPR-barcoding was treated with gefitinib (1 µM) alone or in combination with pemetrexed (50 nM) for 6 days. The proportion of EGFR-T790M cells was assessed by qPCR from genomic DNA and normalized using EGFR_Ctrl primers. Mean ± SEM (n=4) of one representative of three independent experiments. *p < 0.05 (Mann-Whitney test). **C.** Violin plot illustrating the EGFR-score difference between responders (R) and non-responders (NR) from a cohort of renal cell carcinoma patients treated with sorafenib. The EGFR-score was calculated based on the transcriptomic profiles of the tumors before the treatment. The p value was estimated by using the Student t-test. **D.** PC9, HCC4006 or HCC827 NSCLC cells containing EGFR-T790M-mutant subpopulations generated by CRISPR barcoding were treated with gefitinib (1 µM) alone or in combination with sorafenib (5 µM; Soraf) for 5 days and the proportion of EGFR-T790M cells was assessed as in B. Mean ± SEM (n=4) of one representative of at least three independent experiments. *p < 0.05 and **p < 0.01 (Mann-Whitney test). **E.** PC9 and H1975 cells and YU-1150 patient-derived cells (PDC) containing EGFR-C797S-subpopulations were treated with osimertinib (0,1 µM, Osim) alone or in combination with sorafenib (5 µM) for 10 days and the proportion of EGFR-C797S-barcoded cells was measured. Mean ± SEM (n=4) of one representative of at least three independent experiments. **F.** PC9 and H1975 cells described in E were treated with osimertinib alone (1 µM for PC9 cells, 0,1 µM for H1975 cells) or in combination with sorafenib (5 µM) for 20 days (PC9 cells) or 15 days (H1975 cells). The cells were then fixed and stained with crystal violet. **G.** Effects of osimertinib alone (0,1 µM) or in combination with sorafenib (5 µM), sunitinib (1 µM; Sun), regorafenib (2 µM; Reg), lenvatinib (1 µM; Lenv) or cabozantinib (5 µM; Cab) in EGFR-C797S-CRISPR-barcoded PC9 cells. The cells were treated for 7 (left panel) or 10 (right panel) days, and the proportion of the EGFR-C797S subpopulation was measured by qPCR. The mean ± SEM (n=5 or n=4) of one representative of three experiments is shown. **H.** Cell viability of parental or osimertinib-resistant (OR-PC9, EGFR-T790M/C797S) PC9 cells treated for 5 days as indicated with osimertinib (0,1 µM), sorafenib (5 µM), sunitinib (1 µM) or cabozantinib (5 µM). **I.** Colony forming assay representative images of parental, gefitinib-resistant/osimertinib-sensitive (GR-PC9, EGFR-T790M) or osimertinib-resistant PC9 cells treated for 5 days with osimertinib (1 µM) or sorafenib (5 µM), alone or in combination. **J.** Diagram representing the effects of the indicated treatments on a mixed population of EGFR-TKI-sensitive (yellow) and EGFR-TKI-resistant (red) NSCLC cells. **K.** The KRAS-G12D mutation was inserted by CRISPR-barcoding in a subpopulation of H358 cells, and the cells were treated with or without sotorasib (10 nM; Sotor), alone or in combination with sorafenib (5 μM) for 12 days. The proportion of the mutant barcode was assessed by qPCR from genomic DNA and normalized using EGFR_Ctrl primers. Mean ±SEM (n=4) of one representative of three experiments. **L.** Colony forming assay representative images of parental or KRAS-G12D bearing, sotorasib-resistant (SR) H358 cells treated for 8 days with sotorasib (10 nM) or sorafenib (5 µM), alone or in combination.

We confirmed the results of our screen and showed that sorafenib markedly inhibited the gefitinib-induced increase in the proportion of EGFR-T790M mutant cells in different EGFR-addicted NSCLC lines (Figure 1D). Third generation irreversible EGFR-TKIs, such as osimertinib, have been developed to counter resistance induced by the T790M gatekeeper mutation (Cross et al., 2014; Janne et al., 2015). One of the most common mechanisms of resistance to osimertinib in patients is represented by another secondary/tertiary mutation of EGFR, involving the binding site of the drug to the receptor, cysteine 797 (Thress et al., 2015; Passaro et al., 2021). We modeled EGFR-C797S-mediated resistance to osimertinib in PC9 and H1975 cells, as well as in the NSCLC patient-derived cell (PDC) line YU-1150. In each case, co-treatment with sorafenib dramatically reduced the enrichment of EGFR-C797S subpopulations induced by osimertinib, regardless of whether a concurrent T790M mutation was present (Figure 1E-F and Figure S1D).

To assess sorafenib specificity in preventing the emergence of EGFR-TKI resistant NSCLC cells, we tested the effects of other clinically approved multikinase inhibitors displaying partially overlapping pharmacological profiles (Figure S1E). Figure 1G shows that, contrary to sorafenib, sunitinib, regorafenib, lenvatinib and cabozantinib were unable, at commonly used concentrations, to inhibit the enrichment of the EGFR-C797S subpopulations induced by osimertinib.

Next, we derived osimertinib resistant PC9 cells containing the EGFR-C797S mutation (OR-PC9) and compared their response to treatment with that of parental cells. We found that osimertinib inhibited the growth of parental, but not of OR-PC9 cells, whereas sorafenib similarly affected both (Figure 1H-I). As previously shown for pemetrexed, sorafenib didn’t exert any additive/synergistic effects in EGFR-TKI sensitive cells when combined with either osimertinib (Figure 1H-I) or gefitinib (Figure S1F), consistent with the increased vulnerability to sorafenib of cancer cells in which the EGFR pathway is active. These data also explain why this multikinase inhibitor was not considered a hit in previous drug screens aimed at identifying new combinations to prevent resistance in NSCLC cells (Crystal et al., 2014). Together, these results imply that, while EGFR-TKIs alone favor the emergence of resistant over sensitive cells, the combinations of EGFR-TKIs with pemetrexed or sorafenib can inhibit the growth of both cell populations, therefore preserving their relative proportion and preventing tumor evolution (Figure 1J). Cabozantinib, on the contrary, exerts a synergistic effect with osimertinib on sensitive cells, while only poorly affecting resistant cells (Figure 1H), thus resulting in a stronger selective advantage for the latter (Figure J), which explains why this compound, in combination with osimertinib, enhanced the emergence of the subpopulation of EGFR-C797S cells (Figure 1G).

Having demonstrated that the effects of sorafenib on resistant cells cannot be mimicked by other clinically approved multikinase inhibitors, we wanted to investigate whether they are also specific of EGFR-addicted cancer cells. H358 cells derive from human NSCLC and contain the KRAS-G12C driver mutation, which makes them sensitive to a new class of clinically approved KRAS inhibitors, such as sotorasib (Canon et al., 2019). We used CRISPR-barcoding to generate a small subpopulation of cells bearing the KRAS-G12D mutation, a mechanism of resistance to sotorasib identified in patients (Awad et al., 2021). As shown in Figure 1K, treatment with sotorasib induced an enrichment of the proportion of KRAS-G12D cells, which was not affected by co-treatment with sorafenib. Of note, contrary to EGFR-TKIs, we found that these two compounds exert additive effects on cell growth (Figure 1L), implying that, in combination with sotorasib, sorafenib is unable to inhibit the selective advantage of resistant cells. Together, our data indicate that sorafenib can specifically prevent the emergence of EGFR-TKI resistant NSCLC cells.

### Sorafenib prevents the emergence of cells containing different, clinically relevant mechanisms of resistance to EGFR inhibition

Because it targets the capacity of cells to divide, chemotherapy can be effective against different mechanisms of resistance. The results of our initial screen suggested that sorafenib can similarly act on different types of EGFR-TKI-resistant cells. We used CRISPR-barcoding to model other resistance mechanisms found in NSCLC patients (Leonetti et al., 2019; Passaro et al., 2021; Chmielecki et al., 2023). As shown in Figure 2A, co-treatment with sorafenib inhibited the osimertinib-induced enrichment of PC9 subpopulations containing other types of resistance mutations involving receptor tyrosine kinases (RTKs), including EGFR-G724S or an insertion in exon 20 of HER2 (HER2-ex20ins). We then investigated the effects of this drug on cells containing activating mutations of different downstream components of the EGFR signaling. We found that sorafenib prevented the emergence of cells harboring KRAS-G12D, BRAF-V600E or PIK3CA-E545K, in both PC9 cells (Figure 2A) and other NSCLC cell models, including PDCs (Figure 2B and Figure S2A-C). Of note, co treatment with the MEK inhibitor trametinib could inhibit cells containing only some of these mutations (Figure 2A and data not shown).

**Figure 2.**
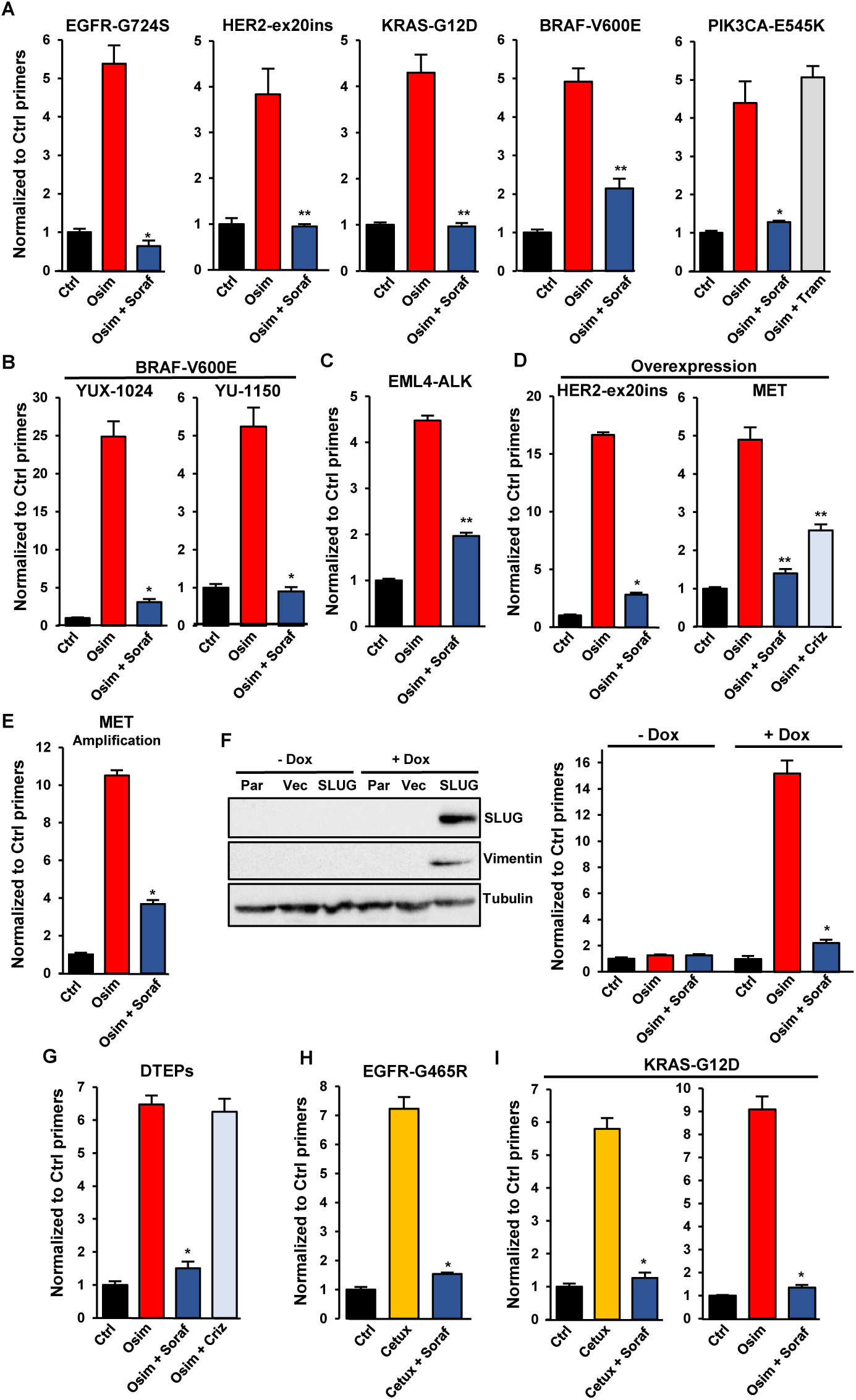
Sorafenib specifically abolishes the selective advantage of cancer cells containing different clinically relevant mechanisms of resistance to EGFR inhibition. **A.** CRISPR-barcoding was used to generate PC9 cell subpopulations containing EGFR-G724S, HER2-ex20ins (HER2-A775insV-G776C), KRAS-G12D, BRAF-V600E or PIK3CA-E545K mutations, and the cells were treated with or without osimertinib (0,1 μM), alone or in combination with sorafenib (5 μM) or trametinib (10 nM; Tram) for 7 (BRAF-V600E), 10 (EGFR-G724S and KRAS-G12D), 15 (PIK3CA-E545K) or 20 (HER-ex20ins) days. The proportions of the mutant barcodes were assessed by qPCR from genomic DNA and normalized using EGFR_Ctrl primers. Mean ±SEM (n=4) of one representative of at least three experiments. *p < 0.05 and **p < 0.01 (Mann-Whitney test). **B.** The BRAF-V600E mutation was introduced by CRISPR/Cas9 in YUX-1024 and YU-1150 PDCs, followed by selection in the presence of osimertinib (0,1 μM). The mutant cells were mixed with parental cells (1 to 100 ratio), and the resulting cell population was treated with osimertinib (0,1 μM, 7 days) alone or in combination with sorafenib (5 μM). The proportion of the BRAF-V600E barcode was measured by qPCR. Mean ±SEM (n=4) of one representative of three experiments. **C.** The EML4-ALK chromosomal inversion was induced by CRISPR/Cas9 in a small subpopulation of PC9 cells. The proportion of mutant cells was assessed after treatment in the presence or the absence of osimertinib (0,1 µM, 10 days), alone or in combination with sorafenib (5 µM). **D.** Overexpression of HER2-A775insYVMA (HER-ex20ins) in PC9 cells was induced by transduction with a lentiviral vector. MET overexpression was induced using a dCas9 activator system through a sgRNA targeting the MET gene, followed by two-week selection in the presence of osimertinib (0,1 μM). The cells overexpressing HER-ex20ins or MET were mixed with parental PC9 cells (1 to 100 ratio) and treated with osimertinib (0,1 μM) alone or in combination with sorafenib (5 μM) or crizotinib (1 µM; Criz) for 10 (HER-ex20ins) or 15 (MET) days. The proportion of cells overexpressing HER-ex20ins or MET was assessed by qPCR from genomic DNA using vector specific primers and normalized using EGFR_Ctrl primers. Mean ±SEM (n=4 for HER2-ex20ins; n=5 for MET) of one representative of three experiments. **E.** EGFR-TKI resistant HCC827-GR6 cells were transduced with a lentiviral vector and mixed with parental HCC827 in a 1 to 100 ratio. The cells were then treated for 6 days with osimertinib (0,1 µM) alone or in combination with sorafenib (5 µM), and the proportion of HCC827-GR6 cells was measured by qPCR using vector specific primers. The mean ±SEM (n=4) of one representative of three different experiments is represented. **F.** PC9 cells were transduced with an empty lentiviral vector (Vec) or a vector for inducible expression of the EMT transcription factor SLUG (SLUG), followed by a 7 day treatment in the presence or the absence of doxycycline (1 µg/ml). Immunoblot (left panel) was performed using anti-SLUG, anti-vimentin and anti-tubulin antibodies. Parental PC9 (Par) were used as negative control. Cells containing the SLUG vector, pre-treated or not with doxycycline for at least one week, were mixed with parental PC9 (1 to 100 ratio) and treated for 7 days with osimertinib (0,1 µM) alone or in combination with sorafenib (5 µM). The proportion of cells containing the lentiviral vector was assessed by qPCR. Mean ±SEM (n=4) of one representative of three experiments. **G.** PC9 cells labeled with a control lentivirus were selected in the presence of osimertinib (0,1 µM) for 3 weeks to obtain a population of drug-tolerant expanded persisters (DTEPs), then mixed with parental cells (1 to 100 ratio). The proportion of DETPS was assessed by qPCR from genomic DNA using primers specific to the lentiviral vector after a 15-day treatment with osimertinib (0,1 µM) alone or in combination with sorafenib (5 µM) or crizotinib (0.5 µM). The mean ±SEM (n=4) of one representative of three experiments is represented. **H-I.** The EGFR-G465R (H) or KRAS-G12D (I) mutations were introduced by CRISPR-barcoding in a subpopulation of LIM1215 CRC cells, and the proportion of mutant cells was assessed after 6 days in the presence or the absence of cetuximab (20 µg; Cetux) or osimertinib (1 µM), alone or in combination with sorafenib (5 μM). The mean ±SEM (n=3 for EGFR-G465R; n=4 for KRAS-G12D) of one representative of three different experiments is represented.

Consistent with our initial screen, sorafenib also blocked the amplification of cells containing the chromosomal inversion leading to the expression of the EML4-ALK fusion oncogene (Figure 2C), suggesting that this combination could also be effective against resistant cells expressing oncogenes originating from RTK fusion, which has been described in a few patients progressing to EGFR-TKIs (Leonetti et al., 2019; Passaro et al., 2021; Chmielecki et al., 2023).

Amplification of other RTKs, including MET and HER2, are among the most common mechanisms of resistance to osimertinib (Leonetti et al., 2019; Passaro et al., 2021; Chmielecki et al., 2023). We modeled these aberrations in PC9 cells using lentivirus mediated cDNA overexpression (Figure S2D) or upregulation of the endogenous gene through a dCas9 activator system (Konermann et al., 2015) (Figure S2E). As shown in Figure 2D, co-treatment with sorafenib strongly inhibited the emergence of cells overexpressing either HER2-ex20ins or MET. To confirm these results, we performed a mixing experiment using parental HCC827 and lentiviral-labeled HCC827-GR6 cells, a MET amplified clone derived by long term selection in the presence of gefitinib (Engelman et al., 2007) (Figure S2F). Figure 2E shows that the enrichment of the HCC827-GR6 subpopulation induced by osimertinib was strongly reduced by co-treatment with sorafenib, indicating that this drug can antagonize acquired resistance mediated by MET amplification.

It has been reported that resistance to EGFR-TKIs can be accompanied by typical features of an epithelial to mesenchymal transition (EMT) (Leonetti et al., 2019; Passaro et al., 2021). Using a lentiviral inducible system, we showed that expression of the EMT transcription factor SLUG can confer resistance to osimertinib in PC9 cells, and this effect was abrogated by co-treatment with sorafenib (Figure 2F). These data imply that this combination can also affect cells that escape EGFR-TKI treatment through phenotypic transformation.

Besides fully resistant cells, it has been shown that certain subpopulations of drug tolerant persisters can survive in the presence of the treatment, in conditions where most of the cells die (Sharma et al., 2010; Hata et al., 2016; Ramirez et al., 2016; Shen et al., 2020). To generate a pool of barcoded drug-tolerant expanded persisters (DTEPs) (Sharma et al., 2010), we treated a mass population of PC9 cells containing a control lentivirus with osimertinib for three weeks. These cells were then mixed with parental PC9 in a 1 to 100 ratio, followed by treatment with osimertinib, alone or in combination with sorafenib or crizotinib. As shown in Figure 2G, osimertinib increased the proportion of DTEPs and this effect was abolished by co-treatment with sorafenib, implying that this multikinase inhibitor can also be effective against non-genetic mechanisms of resistance/tolerance.

Anti-EGFR therapies are used for other tumor types, including colorectal cancers (CRCs) with wt RAS and BRAF, which are treated with anti-EGFR monoclonal antibodies, such as cetuximab or panitumumab. Similarly to NSCLCs, these tumors eventually relapse, due to the acquisition of resistance through various mechanisms, including activation of downstream components of the EGFR pathway or mutations in the extracellular domain of the receptor that prevent its recognition by the antibodies (Di Nicolantonio et al., 2021). We used CRISPR-barcoding to model these mechanisms of resistance in CRC cells sensitive to EGFR inhibition. We found that co-treatment with sorafenib blocked enrichment of CRC cells containing mutations of either EGFR extracellular domain or KRAS induced by cetuximab (Figure 2H-I), indicating that sorafenib could prolong the response to targeted therapy in other types of EGFR-addicted tumors.

### Sorafenib early treatment inhibits MNK activity, STAT3 phosphorylation and MCL1 expression in NSCLC cells

Despite having been originally developed as an inhibitor of the serine/threonine kinase RAF, in PC9 cells sorafenib was unable to down-regulate phospho-ERK levels (Figure 3A) or the expression of well-characterized MAPK-regulated genes, including DUSP6, SPRY2 and ETV5 (Pratilas et al., 2009), as shown by qPCR (Figure S3A). Instead, as previously reported in leukemia cells (Rahmani et al., 2005), sorafenib rapidly inhibited phosphorylation of the eukaryotic translation initiation factor 4E (eIF4E) in PC9 cells (Figure 3A), as well as other NSCLC cells, including two different PDC lines (Figure 3B and Figure S3B), and CRC cells (Figure S3C). A component of the translation regulatory machinery, eIF4E has been implicated in different types of cancer (Ruggero et al., 2004; Wendel et al., 2007; Robichaud et al., 2019; Knight et al., 2021) and its phosphorylation is mediated by MNKs, whose activity is regulated by ERK (Robichaud et al., 2019). Because sorafenib did not detectably affect MAPKs in NSCLC cells and its effects on eIF4E phosphorylation were readily seen after 30 min of treatment (Figure 3A), we hypothesized that this multikinase inhibitor might act directly on MNKs. A potential inhibition of MNK catalytic activity by sorafenib has been suggested by high throughput *in vitro* kinase screens (Davis et al., 2011), although, to our knowledge, this has not been demonstrated in living cells.

**Figure 3.**
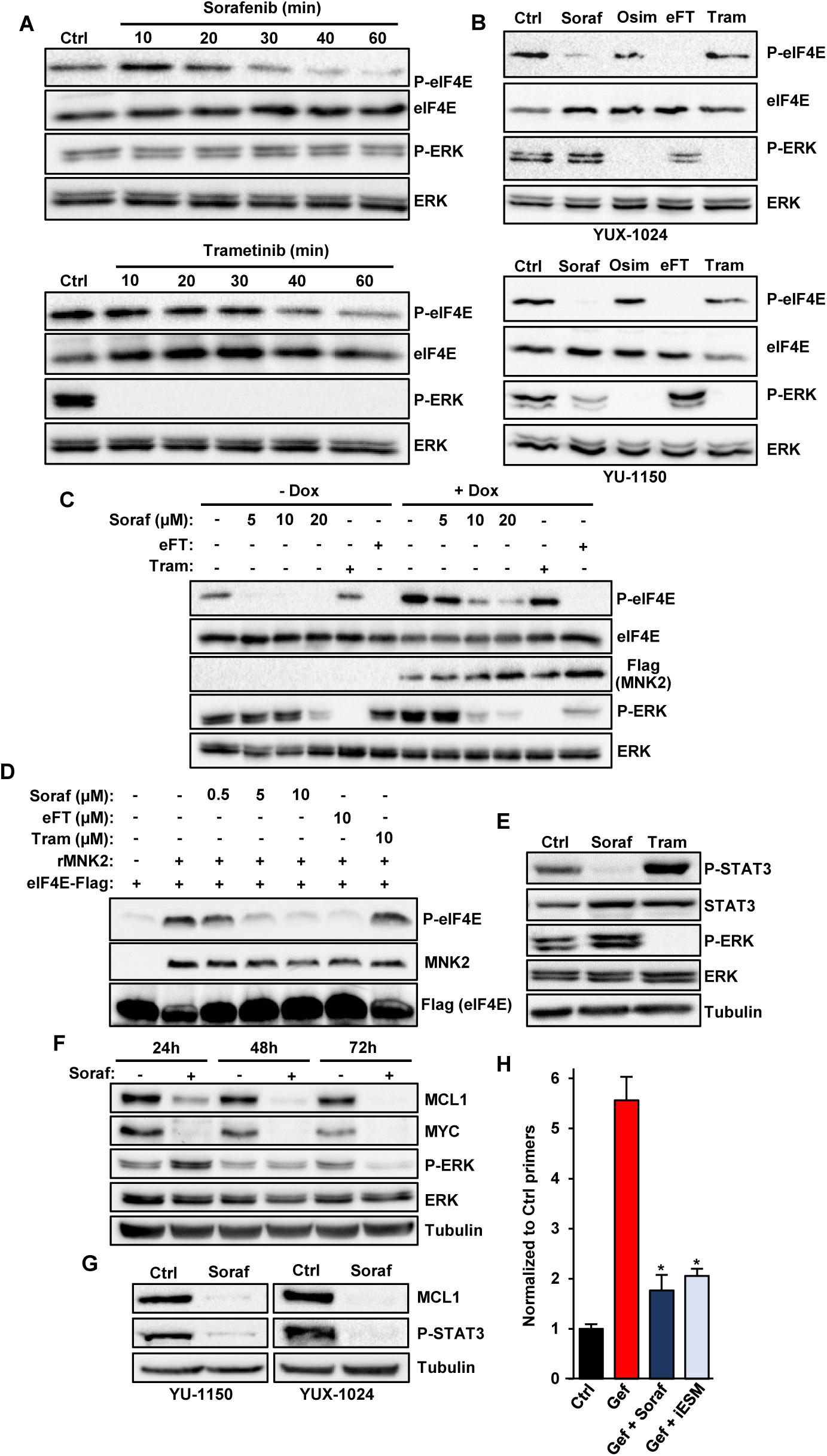
Sorafenib prevents NSCLC cell resistance to EGFR-TKIs independently of MAPKs by inhibiting MNK activity, STAT3 phosphorylation and MCL1 expression. **A.** PC9 cells were treated for 10 to 60 minutes (min) with sorafenib (5 µM) or the MEK inhibitor trametinib (50 nM,), followed by immunoblot using the indicated antibodies. **B.** YUX-1024 and YU-1150 PDCs were treated for 2h with sorafenib (5 µM), osimertinib (0,1 µM), the MNK inhibitor eFT-508 (1 µM; eFT) or trametinib (50 nM; Tram), and immunoblot was performed using the indicated antibodies. **C.** PC9 cells transduced with a lentiviral vector for inducible expression of a constitutively active form of Flag-tagged MNK2 (CA-MNK2) were pre-treated with or without doxycycline (0,1 μg/ml; Dox) for 12h, followed by 1h treatment with sorafenib (5, 10 or 20 µM), eFT-508 (20 μM, positive control) or trametinib (1 μM, negative control). Immunoblot was performed using the indicated antibodies. **D.** Flag-tagged eIF4E was immunoprecipitated from transfected 293T cells and *in vitro* kinase assay was performed using recombinant active MNK2 (rMNK2) in the presence or the absence of the indicated concentrations of sorafenib, trametinib or eFT-508. Immunoblot was performed using anti-phospho-eIF4E, anti-MNK2 and anti-Flag antibodies. **E.** PC9 cells were treated with sorafenib (5 µM) or trametinib (50 nM) for 6h, followed by immunoblot using the indicated antibodies. **F.** PC9 cells were treated with or without sorafenib (5 µM) for 24, 48 or 72h, followed by immunoblot using the indicated antibodies. **G.** YU-1150 and YUX-1024 PDCs were treated with or without sorafenib (5 µM) for 3 days, followed by immunoblot using the indicated antibodies. **H.** EGFR-T790M CRISPR-barcoded PC9 cells were treated with gefitinib (1 µM) alone, or with sorafenib (5 µM) or a combination of the three inhibitors napabucasin (0.5 µM, STAT3 inhibitor), S63845 (0.1 µM, MCL1 inhibitor) and eFT-508 (1 µM; iESM) in 5% FBS medium for 5 days. The proportion of the EGFR-T790M barcode was measured by qPCR from genomic DNA and normalized using EGFR_Ctrl primers. Mean ±SEM (n=5) of one representative of three experiments. *p < 0.05 (Mann-Whitney test).

We initially confirmed that sorafenib can affect MNK1/2 activity in *in vitro* assays, in which the myelin basic protein or a short peptide derived from the human cAMP Response Element Binding protein were used as a substrate (Figure S3D-E). To investigate the potential direct effects of sorafenib on MNK in PC9 cells, we generated a lentiviral inducible system to express a constitutively active form of MNK2 (CA-MNK2). Figure 3C shows that sorafenib, but not the MEK inhibitor trametinib, dose-dependently inhibited the phosphorylation of eIF4E induced by CA-MNK2. In accordance with these results, *in vitro* phosphorylation of immunoprecipitated FLAG-tagged eIF4E by recombinant MNK2 was inhibited by sorafenib, but not by trametinib (Figure 3D). Together, our data demonstrate for the first time that sorafenib can block eIF4E phosphorylation by directly inhibiting MNK catalytic activity. Of note, MNK inhibition by sorafenib didn’t affect cap binding of eIF4E (Figure S3F), suggesting that eIF4E phosphorylation doesn’t exert a global effect on protein translation, consistent with previous studies (Robichaud et al., 2019).

After a few hours of treatment, sorafenib also inhibited the phosphorylation of STAT3 (Figure 3E and Figure S4A), as previously described in other types of cancer (Yang et al., 2008). Of note, the effects on eIF4E and STAT3 phosphorylation were specific for sorafenib and were not observed with other multikinase inhibitors (Figure S4B-C). Sorafenib also induced a more delayed down-regulation of MCL1 and MYC in different NSCLC cells, including two PDC lines (Figure 3F-G and Figure S4D). While previous studies suggested a role of eIF4E phosphorylation in the expression of MCL1 and MYC (Wendel et al., 2007; Knight et al., 2021), the levels of these two proteins were not affected by treatment with the MNK inhibitor eFT508 (data not shown), indicating that inhibition of eIF4E is not sufficient to account for all the effects of sorafenib in NSCLC cells. Of note, while the enrichment of resistant cells could not be blocked by MNK, STAT3 or MCL1 inhibitors alone (Figure S4E and data not shown), the combination of all three compounds mimicked the effects of sorafenib (Figure 3H). Together, these results indicate that sorafenib prevents the emergence of EGFR-TKI resistant subpopulations of NSCLC cells through a mechanism involving, at least in part, its combined inhibitory effects on MNK, STAT3 and MCL1.

### Late down-regulation of EGFR by sorafenib in NSCLC cells

Time-course experiments in PC9 cells revealed that sorafenib also induced a marked down regulation of EGFR expression at the protein level by 2-3 days of treatment, with concomitant inhibition of downstream effectors of the EGFR pathway (Figure 4A and Figure S5A). Similar results were observed with other NSCLC cells, including PDCs (Figure 4B and Figure S5B-C), and CRC cells (Figure S5D), while other multikinase inhibitors, such as sunitinib and regorafenib, failed to effectively down-regulate the receptor (Figure S5E-F). Of note, we didi not observe any effect of sorafenib on EGFR mRNA levels (Figure S5G), implying a post transcriptional effect on protein level. To investigate whether the drug inhibited EGFR protein translation or stability, we generated a lentiviral vector in which the PGK promoter drives expression of FLAG-tagged EGFR and green fluorescent protein (GFP), separated by the T2A self-cleaving peptide. As shown in Figure 4C, sorafenib strongly reduced the expression of the exogenous receptor, whereas it had no effect on GFP levels, indicating that the multikinase inhibitor does not affect EGFR translation, but instead promotes EGFR degradation.

**Figure 4.**
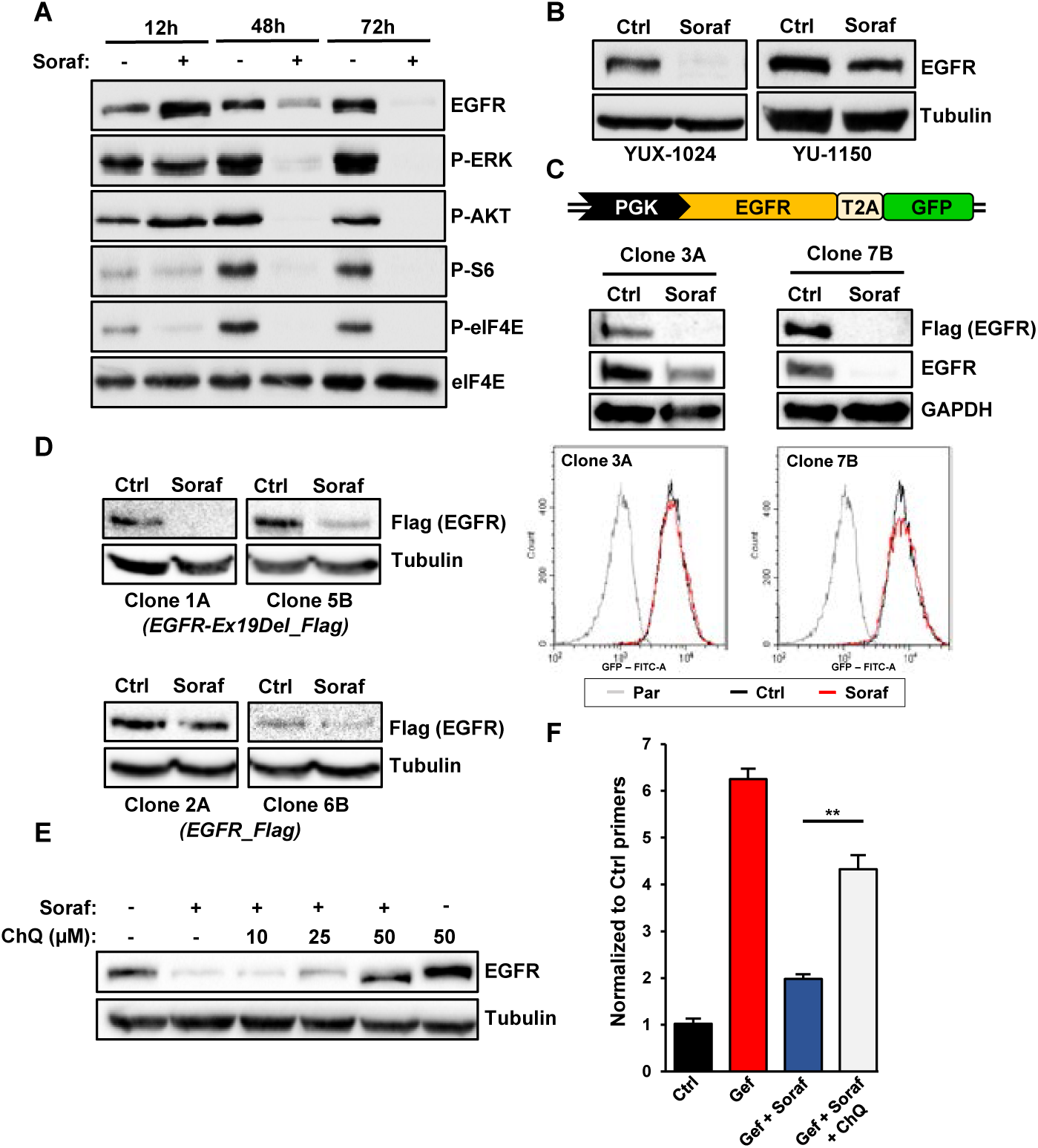
Lysosomal degradation of EGFR participates in the effects of sorafenib in preventing the emergence of EGFR-TKI-resistant NSCLC cells. **A.** Time-course effects of sorafenib (5 µM) in PC9 cells. Immunoblot was performed using the indicated antibodies. Anti-eIF4E was used as a loading control. **B.** YUX-1024 and YU-1150 PDCs were treated with sorafenib (5 µM) for 5 days, followed by immunoblot using anti EGFR and anti-tubulin antibodies. **C.** PC9 cells were transduced with a lentiviral vector containing Flag-tagged EGFR-Ex19Del and the green fluorescent protein (GFP), separated by the T2A self-cleaving peptide and under the control of the PGK promoter. Two different clones were isolated and treated for 3 days in the presence or the absence of sorafenib (5 µM). The expression of EGFR and GFP was measured by immunoblot (upper panel) and FACS (lower panel), respectively. **D.** PC9 cells were transduced with lentiviral vectors containing either Ex19Del-mutant or wt Flag-tagged EGFR, and two clones per condition were isolated. The clones were treated in the presence or the absence of sorafenib (5 µM) for 3 days, and immunoblot was performed using anti-Flag and anti-tubulin antibodies. **E.** PC9 cells were treated with sorafenib (5 µM) alone or in combination with different concentrations of chloroquine (ChQ) for 3 days, followed by immunoblot with anti-EGFR and anti-tubulin antibodies. **F.** PC9 cells containing the EGFR-T790M CRISPR-barcode were treated with gefitinib (1 µM) alone or in combination with sorafenib (5 µM), with or without chloroquine (20 µM) for 4 days. Genomic DNA was derived and the proportion of the EGFR-T790M barcode was measured by qPCR and normalized using the EGFR_Ctrl primers. The mean ±SEM (n=5) of one representative of three experiments is represented. **p < 0.01 (Mann-Whitney test).

It has been reported that, in NSCLC cells, mutant EGFR, but not wt, can be specifically degraded by a macropinocytosis-dependent lysosomal pathway (Menard et al., 2018). Consistent with this mechanism, the inhibitory effects of sorafenib were more pronounced on the Ex19Del mutant compared to the wt receptor (Figure 4D). We also found that suppression of lysosomal degradation by chloroquine blocked the down-regulation of EGFR induced by sorafenib (Figure 4E). Of note, co-treatment with chloroquine also partially rescued the effects of this multikinase inhibitor on the emergence of EGFR-TKI-resistant cells (Figure 4F), implying that its ability to induce EGFR down-regulation contributes to the mechanism of action of sorafenib in NSCLC cells.

### Sorafenib inhibits the clonal evolution of NSCLC cells induced by osimertinib

To further explore the lack of additive effects of osimertinib and sorafenib on growth inhibition, we investigated the effects of these drugs, alone or in combination, on the gene expression profile of PC9 cells (Table S2). As shown by gene set enrichment analysis (GSEA), the profile of osimertinib-treated cells correlated very well with a previously published EGFR-TKI signature (Kobayashi et al., 2006). A similar result was obtained for cells treated with the combination, whereas the enrichment score (ES) was negative in the presence of sorafenib (Figure 5A and Figure S6A), implying that, when sensitive cells are exposed to both drugs, the effects on EGFR-dependent gene expression are predominantly mediated by osimertinib. Consistent with these observations, when we generated our own signature for the osimertinib/sorafenib combination (Table S3), we found higher ES for cells treated with osimertinib, as compared to sorafenib (Figure 5B and Figure S6B). These results were also confirmed by cluster analysis (Figure S6C) and suggest that, in a mixed cell population treated with both drugs, EGFR-TKI sensitive cells respond primarily to osimertinib, whereas sorafenib acts on resistant cells. Mechanistically, we found that the osimertinib/sorafenib combination provoked a decrease of EGFR expression and STAT3 phosphorylation in osimertinib-resistant, but not in osimertinib-sensitive cells (Figure 5C). Similar results were obtained with the gefitinib/sorafenib association (Figure S6D), indicating that, in cells in which EGFR is inhibited, sorafenib is unable to affect phospho-STAT3 and EGFR, which could explain, at least in part, the lack of additive/synergistic effects by the two drugs.

**Figure 5.**
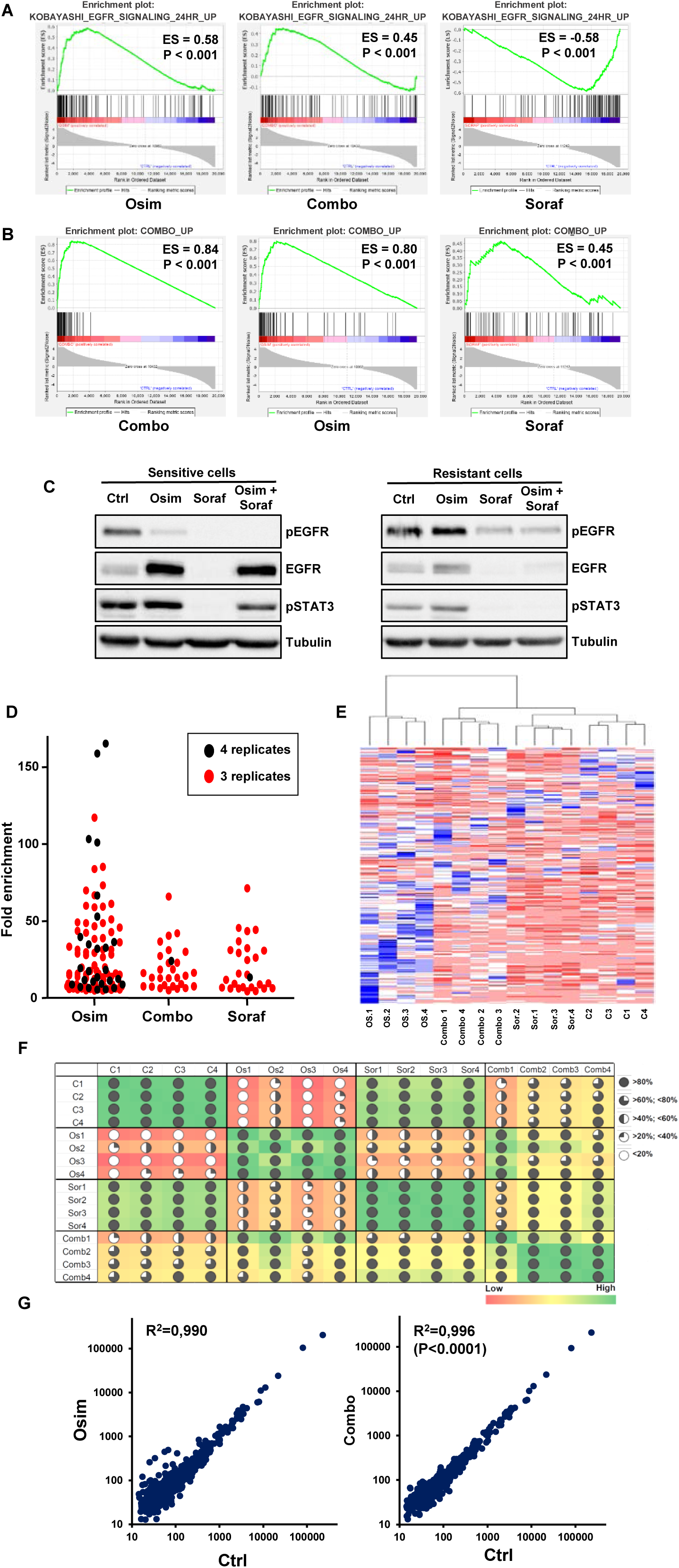
Inhibition of osimertinib-induced clonal evolution by sorafenib in NSCLC cells. **A.** Gene set enrichment analysis (GSEA) of genes up-regulated by EGFR-TKIs in NSCLC cells (KOBAYASHI_EGFR_SIGNALING_24HR_UP signature), performed on gene array data obtained from PC9 cells treated with osimertinib (1 µM) or sorafenib (5 µM), alone or in combination (Combo) for two days. Enrichment scores (ES) and p values are reported. **B.** The data described in **A** were analyzed using our osimertinib-sorafenib combination signature (COMBO_UP). **C.** Osimertinib sensitive and resistant PC9 cells were treated with osimertinib (1 µM) and sorafenib (5 µM), alone or in combination, for 3 days, followed by immunoblot using the indicated antibodies. **D.** PC9 cells containing highly complex CRISPR-barcodes in the AAVS1 locus were treated for two weeks with osimertinib (1 µM) and sorafenib (5 µM), alone or in combination (n=4 per condition). Barcodes enriched at least 5-fold over the control in 4 or 3 replicates are shown. **E.** Barcode heatmap and Ward’s hierarchical clustering of the four replicates per condition of the experiment described in **C**. **F.** Correlation matrix of the data shown in **D**. **G.** Pearson correlation of the barcode distribution in control versus osimertinib or control versus combination treated cells. The coefficients of determination (R^2^) and the p value (Mann-Whitney test) are indicated.

To investigate whether sorafenib could exert a broader inhibition of EGFR-TKI-induced clonal evolution in NSCLC cells, we labeled several thousand PC9 clonal subpopulations using CRISPR highly complex barcodes (Guernet et al., 2016) and compared the effects of a two-week treatment with osimertinib and sorafenib, either alone or in combination. We reasoned that pools of intrinsically tolerant or resistant cells within the original population should be consistently enriched in the different replicates during the treatment, as opposed to randomly selected cells, which would be expected to emerge in only one or two replicates. As shown in Figure 5D and Figure S6E, the number of osimertinib tolerant/resistant subpopulations was strongly reduced by co-treatment with sorafenib, implying a global inhibitory effect on clonal evolution. Consistent with these data, cluster analysis showed that the barcode profile of the cells treated with the combination was closer to that of the control than osimertinib-treated cells (Figure 5E). These results were confirmed by the correlation matrix and correlation plots depicted in Figure 5F and Figure 5G, and indicate that, while osimertinib induces a shift in the clonal architecture of the cancer cell population, this effect can be inhibited by co-treatment with sorafenib.

### The osimertinib/sorafenib combination delays NSCLC resistance *in vivo*

We next sought to investigate the effects of the osimertinib/sorafenib combination *in vivo*. Conventional strategies are generally ill-suited to monitor acquired resistance to this EGFR-TKI in NSCLC mouse models, since the initial response is particularly long, typically lasting for several months (Cross et al., 2014). We thus performed experiments using mass populations constituted of a large majority of osimertinib-sensitive cells and small pools of resistant cells. Using iDISCO 3D imaging (Renier et al., 2014), we showed that EGFR-TKI resistant PC9 cells containing EGFR-C797S, KRAS-G12D or PIK3CA-E545K mutations were enriched in xenografts from mice treated with osimertinib, and this effect was inhibited by co treatment with sorafenib (Figure 6A and Video 1). To better estimate the tumor growth delay induced by this drug combination, SCID mice inoculated with EGFR-C797S CRISPR-barcoded PC9 cells were treated with osimertinib and/or sorafenib. As illustrated in Figure 6B and Figure S7A, the growth of the tumors was slowed by sorafenib and, more markedly, by osimertinib, while it was completely blocked by the combination.

**Figure 6.**
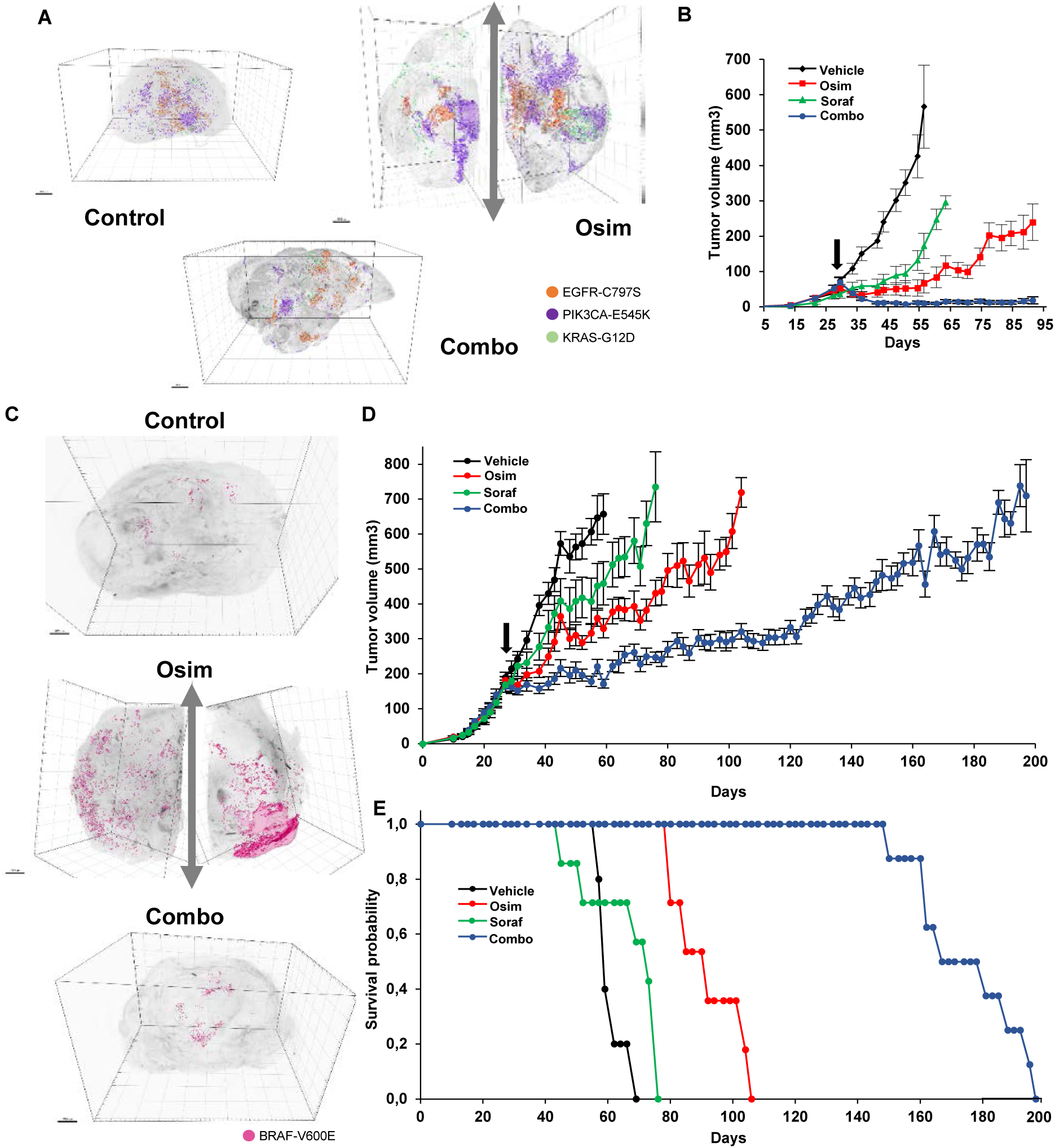
The emergence of subpopulations of osimertinib-resistant NSCLC cells is inhibited *in vivo* by sorafenib. **A.** A PC9 mass population containing small pools (1 to 1000 ratio each) of EGFR-C797S (expressing GFP), KRAS-G12D (expressing β-galactosidase) and PIK3CA-E545K (expressing mCherrry) osimertinib-resistant cells were subcutaneously injected in the right and left flanks of SCID mice. Once the tumors reached a mean volume of about 200 mm^3^, the mice were sacrificed (Ctrl) or treated for 4 weeks with osimertinib (5 mg/kg) alone or in combination with sorafenib (60 mg/kg; Combo). 3D-imaging through iDISCO technology was performed to visualize the subpopulations of resistant cells. To maintain a similar size of the samples from different conditions, the tumors from osimertinib treated mice were cut in two. **B.** A mass population of PC9 cells containing a small pool of cells bearing the EGFR-C797S mutation generated by CRISPR-barcoding was injected in the right and left flanks of SCID mice. Once the tumors were palpable (arrow), the mice were randomized and treated with or without osimertinib (5 mg/kg) and sorafenib (60 mg/kg), alone or in combination, and the volume of the tumors was measured by caliper. The mean tumor volumes ± SEM are represented (n=5 mice per group). **C.** YUX-1024 PDCs containing a subpopulation (1 to 200 ratio) of BRAF-V600E resistant cells expressing GFP were subcutaneously injected in the right and left flanks of SCID mice. Once the tumors reached a mean volume of about 200 mm^3^, the mice were sacrificed (Ctrl) or treated for 4 weeks with osimertinib (10 mg/kg) alone or in combination with sorafenib (60 mg/kg), followed by 3D-imaging. Tumors from osimertinib-treated tumors were cut in two before staining to ensure comparable size of the samples. **D.** A mass population of YUX-1024 PDCs containing a pool of BRAF-V600E cells (1 to 200 ratio) was injected in the right and left flanks of SCID mice. When the tumors reached a mean volume of about 200 mm^3^, the mice were treated with vehicle, osimertinib (10 mg/kg), sorafenib (60 mg/kg) or a combination of the two inhibitors. The mean tumor volumes ± SEM are represented (n=6 mice for the control, n=8 for the other groups). **E.** Kaplan-Meier diagram of the experiment illustrated in **D**. The mice were sacrificed when the volume of at least one of the tumors exceeded 800 mm^3^.

The PDC line YUX-1024 forms tumors that are histopathologically similar to patient derived xenografts obtained from the same patient (Figure S7B). We mixed a small fraction of barcoded BRAF-V600E-mutant cells in a mass population of YUX-1024 cells, which was then inoculated in SCID mice. Osimertinib delayed tumor growth for about a month, while the osimertinib-sorafenib combination was considerably more effective, slowing progression for three additional months (Figure 6C-E and Video 2). Of note, this regimen could be maintained for more than five months without any sign of toxicity.

### Co-treatment with sorafenib prolongs the response to osimertinib in highly aggressive models of resistance in both immunodeficient and immunocompetent mice

The duration of osimertinib response as a single agent is relatively long (Soria et al., 2018a), which makes it more difficult to set up clinical trials to test its effects in combination with other drugs. To investigate whether co-treatment with sorafenib can inhibit the growth of tumors rapidly progressing on osimertinib, we developed a more aggressive model of acquired resistance, which mimics the late phases of osimertinib responsiveness and the beginning of tumor relapse. We generated a PC9 population containing four distinct pools of resistant cells, bearing either EGFR-C797S, PIK3CA-E545K or KRAS-G12D, or overexpressing HER2-ex20ins (Figure S8A). Upon injection in SCID mice, these cells formed tumors whose growth was inhibited by osimertinib by only two weeks, with the rapid amplification of the different subpopulations of barcoded resistant cells (Figure S8B). As shown in Figure 7A-B, co-treatment with sorafenib further delayed tumor growth by more than a month, suggesting that this combination could also be beneficial in patients showing early signs of progression to osimertinib.

**Figure 7.**
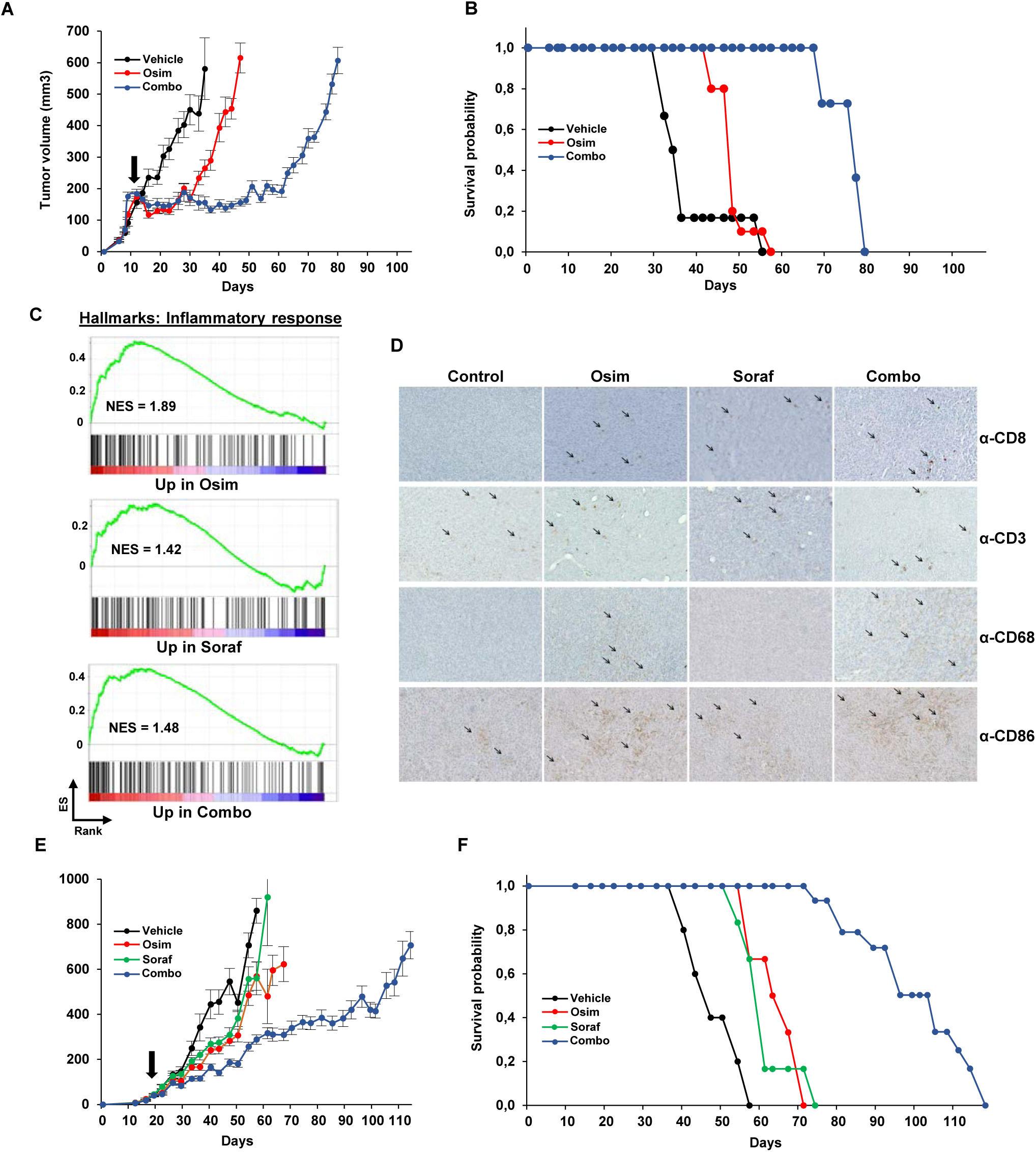
The osimertinib-sorafenib combination promotes the recruitment of antitumor immune cells and substantially prolongs the response in extremely aggressive models of acquired resistance. **A.** A PC9 mass population containing small pools of EGFR-C797S, KRAS-G12D and PIK3CA-E545K CRISPR-barcoded or HER2-ex20ins overexpressing cells was injected in the right and left flanks of SCID mice. Once the tumors reached a mean volume of about 100 mm^3^, the mice were randomized and treated with either vehicle or osimertinib (5 mg/kg), alone or in combination with sorafenib (60 mg/kg) and the volume of the tumors was measured by caliper. The mean tumor volumes ± SEM are shown (n=6 mice for the control, n=10 mice for the other groups). **B.** Kaplan-Meier diagram of the experiment shown in **A**. The mice were sacrificed when the volume of at least one of the tumors exceeded 800 mm^3^. **C.** GSEA of the genes up-regulated by osimertinib, sorafenib or the combination in PC9 cells based on an inflammatory response signature. **D.** Mouse BEM-5 cells addicted to mutant EGFR were injected in the right and left flanks of syngeneic BALB/c mice. When the tumors reached a mean volume of about 50 mm^3^, the mice were treated for 10 days with vehicle, osimertinib (20 mg/kg), sorafenib (60 mg/kg) or a combination of the two inhibitors, followed by IHC analysis using the indicated antibodies. Images representative of three different tumors/mice per condition are shown. **E.** BEM-5 cells were injected in the right and left flanks of BALB/c mice. Once the tumors reached a mean volume of about 50 mm^3^, the mice were randomized and treated with either vehicle or osimertinib (20 mg/kg) alone or in combination with sorafenib (60 mg/kg). The mean tumor volumes ± SEM are shown (n=5 mice for control, n=6 mice for osimertinib and sorafenib, n=16 mice for the combination). **F.** Kaplan-Meier diagram of the experiment shown in **E.** The mice were sacrificed when the volume of at least one of the tumors exceeded 800 mm^3^.

EGFR-mutant NSCLCs have been reported to respond poorly to anti-PD1/PDL1 inhibitors (Gainor et al., 2016). However, it has been shown that EGFR signaling can promote a noninflamed tumor microenvironment, and inhibition of this receptor can stimulate the infiltration of inflammatory immune cells (Akbay et al., 2013; Ayeni et al., 2019; Sugiyama et al., 2020). These observations suggest that acquisition of EGFR-TKI resistance could be influenced by the presence of a functional immune system. We first tested whether treatment of NSCLC cells with osimertinib or sorafenib could affect the expression of immune-related genes. As shown in Figure 7C, GSEA revealed up-regulation of an inflammatory response signature in PC9 cells treated with both inhibitors, either alone or in combination, suggesting that these drugs might enhance tumor recognition by the immune system. Transgenic models are poorly suited to investigate acquired resistance and the few available mouse NSCLC lines don’t contain mutant EGFR.

To investigate whether sorafenib might prolong the response to osimertinib in immunocompetent mice, we generated a new syngeneic model of oncogenic addiction to mutant-EGFR by transforming BALB-3T3 cells (Aaronson and Todaro, 1968) using a lentiviral vector containing mouse Egfr-L860R (corresponding to human EGFR-L858R). Compared to cells expressing GFP or a dominant-negative p53 (DN-p53), BALB-3T3-Egfr-L860R cells are highly sensitive to osimertinib (Figure S9A). Consistent with these data, these cells can be made unresponsive to EGFR-TKIs upon insertion of a resistance mutation by CRISPR-barcoding (Figure S9B). As shown in Figure S9C, BALB-3T3-Egfr-L860R cells formed rapidly growing tumors in BALB/c mice. To obtain a homogeneous population of tumorigenic cells, we dissected the tumors and isolated several clones sensitive to osimertinib from the disaggregated cells (Figure S9D). We then took one of these clones (BALB-3T3-Egfr-L860R-M5, hereafter BEM-5 cells) and generated by CRISPR-barcoding a subpopulation of osimertinib resistant cells containing the mouse Egfr-C799S mutation (corresponding to EGFR-C797S in the human receptor). Upon injection in immunocompetent mice, these cells formed aggressive tumors that showed short-term response to osimertinib. To investigate the effects of treatment on the recruitment of immune cells, once the tumors reached a volume of about 50 mm^3^ the mice were treated for 10 days in the presence or the absence of osimertinib and sorafenib, alone or in combination, followed by immunohistochemistry analysis. As shown in Figure 7D and S9E, treatment with the inhibitors increased the number of CD8 positive cells in the tumors, while osimertinib and the osimertinib/sorafenib combination also enhanced the recruitment of proinflammatory macrophages, consistent with an immunosuppressive effect of EGFR signaling. We then compared the effects of these inhibitors on the growth of BEM-5 tumors in immunocompetent mice. While osimertinib and sorafenib alone slowed the growth of the tumors by about two weeks compared to the control, the combination induced an additional delay of almost two-months (Figure 7E-F). Together, our results indicate that co-treatment with sorafenib can substantially prolong the therapeutic response to osimertinib even in rapidly progressing tumors, and this effect may be enhanced by activation of an antitumor response by the immune system.

## DISCUSSION

Clinically approved for two decades to treat NSCLC, EGFR-TKIs typically show high response rates and prolonged efficacy compared to other targeted therapies. Unfortunately, despite the strong addiction of NSCLCs to mutant EGFR and the high selectivity of these drugs for the mutant form of the receptor, these tumors almost inexorably relapse and become resistant to this treatment. To prevent this process, various strategies have been proposed to either achieve deeper inhibition of EGFR signaling, or target a specific mechanism of tolerance/resistance (Engelman et al., 2007; Cross et al., 2014; Tricker et al., 2015; Lee et al., 2018; Shah et al., 2019; Kurppa et al., 2020; Tanaka et al., 2021; Noronha et al., 2022; To et al., 2022). However, complete inhibition of the pathway can cause unacceptable toxicity to normal tissues, and multiple mechanisms of tolerance/resistance generally coexist in the same patient. While acquired resistance to EGFR-TKIs in advanced NSCLC may be ineluctable, recent clinical trials have shown that co-treatment with chemotherapy can significantly delay progression (Hosomi et al., 2020; Noronha et al., 2020).

Our present findings establish that chemotherapy functions, at least in part, by inhibiting the growth of EGFR-TKI resistant cells. Thus, these cells can be more vulnerable towards certain types of drugs because of their capacity to proliferate in the presence of EGFR-TKIs. By screening for other compounds that might be effective in inhibiting the emergence of cells with different mechanisms of resistance to EGFR-TKIs, we identified sorafenib and showed it to be specific for EGFR addicted cancer cells. Moreover, its effects could not be mimicked by other multikinase inhibitors. Importantly, though initially developed as a RAF inhibitor, we observed that its effects were mediated through several other intracellular pathways rather than through rapid inhibition of RAF signaling. We showed for the first time that sorafenib rapidly blocked the catalytic activity of MNKs, which promote tumor progression by phosphorylating eIF4E, associated with efficient translation of several oncogenes (Wendel et al., 2007; Knight et al., 2021), and through other mechanisms as well (Brown and Gromeier, 2017). At later time points, we showed that sorafenib inhibits STAT3 phosphorylation, as well as the expression of MCL1, MYC and EGFR. While MNK and MYC inhibition were induced by both sorafenib and EGFR-TKIs (Figure 3 and data not shown), decreased levels of EGFR and phospho-STAT3 were observed in the presence of sorafenib alone, but not with the combination, hence providing a mechanistic basis for the lack of synergy between the two drugs. Together, our findings imply that, in a population of NSCLC cells treated with the combination, osimertinib acts on EGFR-TKI-sensitive cells, whereas sorafenib inhibits the growth of EGFR-TKI-resistant cells (Figure S10). Thus, by relieving the selective pressure induced by osimertinib, sorafenib counteracts the capacity of the tumor cell population to adapt through clonal evolution.

Besides its approval for liver, thyroid and kidney cancer, sorafenib has been tested in other malignancies, including lung cancer. From a cohort of heavily pretreated patients, the BATTLE trial reported higher eight-week disease control rate for tumors containing wt EGFR. However, no significant differences in progression free-survival (PFS) were observed, and the study included only 12 patients with mutant EGFR (Kim et al., 2011; Blumenschein et al., 2013). The opposite conclusion was drawn from a larger, phase III trial involving 89 and 258 patients displaying mutant and wt EGFR, respectively. Indeed, in the placebo-controlled MISSION study, sorafenib significantly improved both PFS (2.7 months for sorafenib versus 1.4 months for placebo) and OS (13.9 versus 6.5 months) in NSCLC patients with mutant EGFR, whereas no difference was observed when the whole cohort of patients was analyzed (Paz-Ares et al., 2015). These data are consistent with our EGFR-score analysis and indicate that NSCLCs with activation of this receptor are more sensitive to sorafenib.

A few studies have also tested the effects of sorafenib in combination with a first generation EGFR-TKI in NSCLC patients. While some of these were only designed to evaluate the tolerability of the treatment (Adjei et al., 2007; Duran et al., 2007), three multicenter phase II trials were performed to investigate the efficacy of the erlotinib-sorafenib combination. Two of these studies did not compare the effects of the combination with those of erlotinib alone, and the authors reported a better response in the few patients with mutant EGFR tumors (Lind et al., 2010; Lim et al., 2016). The third trial found no difference between the combination and the EGFR-TKI alone, but it only involved five patients with mutant EGFR (two in the combination and three in the erlotinib group) (Spigel et al., 2011). Similarly to the early clinical trials that failed to prove the efficacy of the chemotherapy-EGFR-TKI association (Rotow and Janne, 2020), the very limited number of EGFR-mutant tumors in these cohorts preclude any meaningful evaluation of the potential benefit of a sorafenib-EGFR-TKI combination. However, all these studies reported that the treatment was generally well-tolerated, suggesting that the adverse effects of the association of osimertinib with sorafenib should also be manageable.

To clinically test a new drug combination, the expected benefits for the patients need to counterbalance the increased toxicity. Because the median progression-free survival of osimertinib as a single agent exceeds 18 months (Soria et al., 2018b), the set up of new clinical trials in treatment-naïve NSCLC patients can pose ethical concerns. To investigate the potential effects of our combination on more advanced tumors, we generated mouse models that mimicked rapid progression to EGFR-TKI treatment. We showed that co treatment with sorafenib can substantially prolong the response of these aggressive tumors to osimertinib, indicating that the combination could also be beneficial during later phases of the disease.

Using a new syngeneic model of oncogenic addiction to mutant EGFR, we showed that the osimertinib-sorafenib combination is also effective in immunocompetent mice, where it promoted an inflammatory response by the immune system. These findings are consistent with the immunosuppressive effects of EGFR signaling in NSCLC described in previous studies (Akbay et al., 2013; Ayeni et al., 2019; Sugiyama et al., 2020) and imply that the immune system could play a role in delaying the acquisition of osimertinib resistance. A contribution by the immune system might also explain the higher degree on variability in the individual response to the combination observed in immunocompetent versus immunodeficient mice (Kaplan-Meir diagrams of Figure 7) and suggest that an analysis of the levels of infiltrated inflammatory cells induced by osimertinib may help to predict the duration of the response.

In conclusion, we showed that, similarly to chemotherapy, sorafenib can delay the acquisition of NSCLC resistance to EGFR-TKIs. While less efficient than EGFR-TKIs in inhibiting the growth of EGFR mutant cancer cells, sorafenib and chemotherapy specifically target the emerging subpopulations of resistant cells, thus prolonging NSCLC response to EGFR-TKIs. According to the toxicities observed in the early clinical trials with first generation EGFR-TKIs described above (Lind et al., 2010; Spigel et al., 2011; Lim et al., 2016;Hosomi et al., 2020; Noronha et al., 2020), sorafenib is expected to be more tolerable than chemotherapy in combination with osimertinib. Moreover, sorafenib can also affect tumor angiogenesis through VEGFR and PDGFR inhibition, and different studies have shown that co-treatment of EGFR-TKIs with antiangiogenic drugs can improve progression free survival of NSCLC patients (Nakagawa et al., 2019; Saito et al., 2019; Zhou et al., 2021). Altogether, our findings strongly support the clinical potential of combined osimertinib and sorafenib therapy for EGFR mutant NSCLCs.

## METHODS

### Cell culture and Inhibitors

Human embryonic kidney 293T cells were obtained from ATCC; PC9 cells (NSCLC, EGFR-Ex19Del) were obtained from ECACC (distributed by Sigma-Aldrich); HCC4006 (NSCLC, EGFR-Ex19Del) and H1975 cells (NSCLC, EGFR-L858R/T790M) were obtained from ATCC; HCC827 cells (NSCLC, EGFR-Ex19Del) and H358 (NSCLC, KRAS-G12C) were a gift from Pr. J. Minna, UT Southwestern Medical Center, Dallas (USA); HCC827-GR6 were a gift from Pr. P. Jänne, Dana-Farber Cancer Institute, Boston (USA); HCA-46 cells (CRC) were obtained from ECACC; CCK81 cells (CRC) were obtained from HSRRB (Japan); LIM1215 cells (Whitehead et al., 1985) were obtained from Prof. R. Whitehead, Vanderbilt University, Nashville (USA), with permission from the Ludwig Institute for Cancer Research, Zurich (Switzerland). NSCLC PDC lines YU-1150 (EGFR-L858R/T790M) and YUX-1024 (EGFR-L858R) were described previously (Yun et al., 2019). BALB-3T3 cells were derived from BALB/c embryos (Aaronson and Todaro, 1968). The different cell lines were authenticated by short tandem repeat profiling (Eurofins Genomics). 293T cells were grown in DMEM (Life Technologies) supplemented with 10% FBS (Life Technologies); HCA-46 cells were grown in DMEM supplemented with 5% FBS; PC9, HCC4006, HCC827, H1975, YU-1150 and YUX-1024 were grown in RPMI1640 (2Mm L-Glutamine + 25 mM HEPES) supplemented with 10% FBS; LIM1215 cells were grown in RPMI1640 supplemented with 10% FBS, 0.5 µg/ml insuline (Sigma-Aldrich), 1 µg/ml hydrocortisone (Sigma-Aldrich) and 10 µM 1-thioglycerol (Sigma-Aldrich); CCK81 cells were grown in MEM (Life Technologies) supplemented with 10% FBS. BALB-3T3 cell were grown in DMEM supplemented with 10% calf serum (Life Technologies). All media were supplemented with 0.5% penicillin/streptomycin (Life Technologies).

Osimertinib, sorafenib, cabozantinib and lenvatinib were purchased from LC Laboratories. Gefitinib and doxycycline were purchased from Santa Cruz Biotechnology. Trametinib and puromycin were purchased from ChemCruz. Pemetrexed, sunitinib, eFT-508, napabucasin, S63845, cetuximab were purchased from Selleckchem. Regorafenib was purchased from TargetMol. Crizotinib and chloroquine were purchased from Sigma-Aldrich. Sotorasib was purchased by MedChemExpress. Blasticidin and zeocin were purchased from ThermoFisher.

### CRISPR-Barcoding, cell transfection and DNA constructs

sgRNA vectors were generated as previous described (Guernet et al., 2016) by cloning annealed oligonucleotides containing the targeting sequence (Table S4) into the pSpCas9(BB)-2A-Puro vector (a gift from Feng Zhang, Addgene plasmid # 48139) (Ran et al., 2013) digested with BbsI (New England Biolabs). The sequences of the donor single stranded DNA oligonucleotide (ssODNs, Integrated DNA technologies) used for CRISPR/Cas9-mediated homology directed repair, containing the mutation of interest and a few additional silent mutations constituting the barcode, are shown in Table S5. Cells were co-transfected with 2 µg of the CRISPR/Cas9 plasmid and 2 µg of ssODN (50 µM) using a Nucleofector II and Amaxa Nucleofector kits (Lonza) according to the manufacturer’s instructions. The efficiency of each transfection was checked in parallel using a GFP containing plasmid.

Lentiviral vectors for inducible expression of CA-MNK2 (T379D) and SLUG were generated by Gibson assembly (New England Biolabs) or standard cloning in the pTRIPZ plasmid from pTK-Slug (a gift from Robert Weinberg, Addgene plasmid # 36986) (Guo et al., 2012) and pDONR223-MKNK2 (a gift from William Hahn & David Root, Addgene plasmid # 23735) (Johannessen et al., 2010).

The sequence encoding the Flag tag was cloned between the HindIII and KpnI sites of the pHA-eIF4E plasmid (a gift from Dong-Er Zhang, Addgene plasmid # 17343) (Okumura et al., 2007). Flag-tagged wt and Ex19Del human EGFR constructs in the lentiviral VIRSP vector were generated by PCR with Herculase II Fusion DNA Polymerase (Agilent technologies) from EGFR WT (a gift from Matthew Meyerson, Addgene plasmid # 11011) (Greulich et al., 2005) and pBabe EGFR Del1 (a gift from Matthew Meyerson, Addgene plasmid # 32062) (Yuza et al., 2007) plasmids, respectively. The T2A peptide and EGFP were cloned by Gibson assembly downstream of EGFR-Ex19Del in the VIRSP plasmid. WT HER2 was cloned into VIRSP from the pCEV29-erbB2 plasmid previously generated by Makoto Igarashi in the Aaronson lab, and the sequence encoding A775insYVMA was added by site-directed mutagenesis and Gibson assembly. Mouse EGFR-L860R (corresponding to human EGFR-L858R) was clone into VIRSP by PCR amplification from cDNA derived from a mouse hypothalamic cell line (Cellutions Biosystems) and Gibson assembly. For MET overexpression through CRISPRa (Konermann et al., 2015), the lentiviral vectors dCAS-VP64_Blast, MS2-P65-HSF1_Hygro and sgRNA(MS2)_zeo (gifts from Feng Zhang, Addgene plasmids # 61425, # 61426 and # 61427) were used. The MET sgRNA target sequence (5’-TGGCAGGGCAGCGCGCGTGT) was cloned in sgRNA(MS2)_zeo. To generate osimertinib-tolerant cells, PC9 cells transduced with dCAS-VP64_Blast, MS2-P65-HSF1_Hygro and empty sgRNA(MS2)_zeo were used. To label cells used in iDISCO 3D-imaging, lentiviral vectors GFP-VIRHD, mCherry-VIRHD and pLenti-puro-LacZ (a gift from Ie Ming Shih, Addgene plasmid # 39477) (Guan et al., 2011) were used. All constructs were sequence verified.

For lentivirus production, 293T cells were co-transfected using polyethylenimine (Polysciences) with the lentiviral vector, pCMV Δ8.91 and pMD VSV-G plasmids as previously described (Grumolato et al., 2010). The conditioned media containing the viral particles was collected two, three and four days after transfection, cleared by centrifugation, supplemented with 8 µg/ml polybrene and added to PC9 cells for overnight incubation at 37°C, 5% CO_2_. Two days after transduction, the cells were selected in 1 µg/ml puromycin, 10 µg/ml blasticidin or 250 µg/ml zeocin.

### DNA extraction, RNA extraction and qPCR

Genomic DNA (gDNA) was extracted using NucleoSpin Tissue Kit (Macherey-Nagel) according to the manufacturer’s instructions. Total RNA was isolated using the Tri-Reagent (Sigma-Aldrich) and chloroform (Sigma-Aldrich), purified with NucleoSpin RNA columns (Macherey-Nagel), quantified by Nanodrop One (Thermo Scientific), DNase digested and reverse-transcribed using the Improm-II Reverse Transcription System (Promega). qPCR was performed from 100 ng of gDNA using the Fast SYBR Green Master Mix (Applied Biosystems) on a QuantStudio Flex PCR System (Thermo Scientific). The sequence of the different PCR primers, designed using Primer-BLAST (NCBI), is provided in Table S6. For CRISPR-barcoding experiments, to avoid potential amplification from ssODN molecules not integrated in the correct genomic locus, one of the two primers was designed to target the endogeneous genomic sequence flanking the region sharing homology with the ssODNs. qPCR analysis was performed using the standard curve method. For the initial CRISPR-barcoding screen, qPCR was performed using a QuantStudio 12K Flex System (Thermo Scientific). The wt and mutant EGFR-T790 barcodes were detected by Taqman PCR using the EGFR-FW (5’-TCCCTCCAGGAAGCCTACG) and EGFR-RV (5’-CCTTCCCTGATTACCTTTGCGA) primers and the EGFR-T790T (5’-CCAACTGATTACCCAGCTCATGCCC) and the EGFR-T790M (5’-GCTTATAATGCAACTGATGCCCTTCGG) probes, using a Taqman master mix (Thermo Scientific).

### Cell viability and colony forming assays

For cell viability assay, 2,500 PC9 cells per well were seeded in 96-well plates. Cells were incubated overnight to allow attachment to the plates, followed by treatment with the different drugs. The culture media/drugs were refreshed every 2/3 days. After 6 days of treatment, CellTiter-Glo reagent (Promega) was added to culture medium, mixed with an orbital shaker for 2 min to induce cell lysis and incubated for 10 min at room temperature to stabilize the luminescent signal. Measurements were performed according to manufacturer’s instructions using Infinite F200 PRO (TECAN). Cell viability was analyzed based on the levels of luminescence, proportional to the amount of ATP present in the cells.

For long-term colony forming assays, 5,000-10,000 cells per well were seeded in six well plates. The following day, the cells were treated as indicated for the different experiments, then they were dried, fixed with a 10% MetOH, 10% acetic acid solution, and stained with a 1% crystal violet (Merck) MetOH solution. Excess stain was removed by rinsing in deionized water and the plates were air-dried. Plate images were captured using a ChemiDoc Imaging System.

### EGFR-score analysis

The EGFR-score reflects activation of EGFR signaling and was calculated as described previously (Cheng et al., 2020) based on the gene expression data (GSE180925) from a cohort of renal cell carcinomas treated with sorafenib (Gudkov et al., 2022).

### Flow cytometry

Parental or Flag-EGFR_Ex19Del-T2A-GFP (VIRSP)-transduced PC9 cells, treated with or without sorafenib (5 µM) for three days, were trypsinized, washed and fixed with 4% paraformaldehyde. The levels of green fluorescence were measured by FACS using a CytoFLEX Flow Cytometer (Beckman Coulter) and analysed with the CytExpert software (Beckman Coulter).

### Immunoblot, m7GTP pull-down and *in vitro* kinase assay

Cells were lysed in a buffer containing 50 mM HEPES pH 7.6, 150 mM NaCl, 5 mM EDTA, NP40 0.5%, 20 mM NaF, 2 mM Na3VO4, supplemented with protease inhibitor mini tablets (Thermo Scientific). Lysates were cleared by centrifugation at 14,000 g for 15 min at 4°C and protein concentration was determined using the Bradford assay (Bio-Rad). Sodium dodecyl sulfate (SDS) loading buffer was added to equal amounts of lysates, followed by SDS polyacrylamide gel electrophoresis (PAGE) and transfer to polyvinylidene fluoride membranes (Bio-Rad) using a Trans-Blot Turbo Transfer System (Bio-Rad). Membranes were blocked for 1 h at room temperature in 5% nonfat dried milk (Sigma-Aldrich) in PBS and incubated overnight at 4°C with the primary antibodies (Table S7) in 3% BSA PBS-tween. After washing, the membranes were incubated 1 h with horseradish peroxidase-conjugated secondary antibodies (1:5,000) and the bands were visualized by chemiluminescence using the Clarity Western ECL substrate (Bio-Rad) and a ChemiDoc Imaging System (Bio-Rad). The images were analyzed using the Image Lab Software (Bio-Rad).

To isolate cap-binding proteins, 40 µL of immobilized gamma-aminophenyl-m7GTP beads (Jena Bioscience) were added to 1 mg of PC9 cell lysates, followed by overnight incubation at 4°C. After four washes in lysis buffer, SDS loading buffer was added to the beads, and immunoblot was performed in parallel from both the m7GTP-bound fraction and total lysate.

For *in vitro* MNK2 kinase assay, 293T cells transfected with Flag-eIF4E (pcDNA3) were treated overnight with trametinib 0.1 µM, followed by cell lysis and immunoprecipitation using anti-Flag M2 agarose beads (Sigma-Aldrich). After extensive washing with cold lysis buffer and PBS, equal amounts of beads were incubated with kinase buffer (Abcam) supplemented with 300 µM ATP, 0.25 mM DTT and 50 µg/ml BSA, in the presence or the absence of 100 ng of recombinant human MNK2 protein (Abcam) and different concentrations of sorafenib or other inhibitors for 20 min at 30°C. The reaction was stopped by adding SDS loading buffer, followed by SDS-PAGE and immunoblot.

### Gene array analysis

PC9 cells were treated in triplicate with or without osimertinib (1 µM) and/or sorafenib (5 µM) for 1, 2 or 4 days (for the longest time point, the media was renewed after 2 days), and total RNA was isolated using the miRNeasy kit (Qiagen) following the manufacturer’s instructions. The RNA quality was verified on agarose gel. Gene expression profiling was performed using a low-input QuickAmp labeling kit and human SurePrint G3 8×60K microarrays (Agilent Technologies), as previously described (Sulpice et al., 2016). Differentially expressed genes were identified by a 2-sample univariate t test and a random variance model, as previously described (Allain et al., 2016).

### Barcode sequencing

Highly complex CRISPR-barcodes were inserted into the AAVS1 locus of PC9 cells as previously described (Guernet et al., 2016). Barcoded cells were treated for 2 weeks with or without 1 μM osimertinib and 5 μM sorafenib, alone or in combination (four replicates per condition, one million cells per replicate in T-150 cm^2^ flasks). After treatment, the cells were harvested and genomic DNA was extracted using NucleoSpin® Tissue kit (Macherey-Nagel) according to manufacturer’s instructions. Targeted amplification of the integrated barcode in the AAVS1 locus was performed using previously described (Guernet et al., 2016) primers containing Illumina adapter sequences and 6 bp unique indexes. For each sample, we performed 3 PCR reactions, each from 500 ng of genomic DNA in a final volume of 50 µl, using Herculase II Fusion DNA Polymerase (Agilent technologies) and the following program: 98°C for 5 minutes; followed by 27 cycles of 20 s at 98°C, 20 s at 60°C and 30 s at 72°C; final extension at 72°C for 3 min. The PCR products from the same sample were pooled and purified over 2% agarose gels (band size at 275 bp) using NucleoSpin® Gel and PCR clean up kit (Macherey-Nagel). Purified amplicons were quantified by Qubit (Thermofisher) and their quality was assessed by Bioanalyzer (Agilent). Sequencing was performed on an Illumina MiniSeq, using the High Output Reagent Kit (300-cycles). Counts of barcodes for each sample were extracted from FASTQ files using galaxy (https://usegalaxy.org). Heatmap and Ward’s clustering analyses were performed using euclidean distance with R software. Spearman correlations were calculated using Excel (Microsoft).

### Generation of a syngeneic model of oncogenic addiction to mutant EGFR

BALB-3T3 cells were transduced with the lentiviral mouse EGFR-L860R vector and selected for 5 days in the presence of puromycin. To enrich for cells with stronger EGFR signaling activation, the cells were serum starved over two cycles of two weeks. Addiction to EGFR-L860R was assessed by colony forming assay in the presence of different concentrations of osimertinib. To select for tumorigenic cells, BALB-3T3-EGFR-L860R cells were bilaterally inoculated in the flanks of BALB/c mice. The tumors were dissociated and the cells grown in culture for one week in the presence of puromycin to eliminate the remaining cells from the host. Several clones were derived from the mass population by limiting dilution, and their reliance on EGFR signaling was assessed by colony forming assay (data not shown) and immunoblot.

### Mouse xenografts

For the experiment illustrated in Figure 6B, PC9 cells containing the EGFR-C797S CRISPR-barcode were mixed with Matrigel (Corning) and subcutaneously inoculated in the left and right flanks (2 × 10^6^ cells per site) of male SCID mice. For the experiment shown in Figure 7A-B, separate batches of PC9 cells containing the EGFR-C797S, KRAS-G12D or PIK3CA-E545K CRISPR-barcodes were pooled and supplemented with a small fraction (1:200) of PC9 cells transduced with the HER2-ex20ins vector. The cells were mixed with Matrigel and bilaterally inoculated in male and female SCID mice. For the experiment depicted in Figure 6D-E, YUX-1024 cells were transfected for BRAF-V600E CRISPR-barcoding and selected with 0.1 µM osimertinib for three weeks. The cells were then mixed with parental YUX-1024 in a 1:200 proportion, supplemented with 50% Matrigel and bilaterally inoculated in the flanks of female SCID mice. The size of the tumors was measured by caliper every 2-3 days. When tumors were palpable (Fig 6B) or reached an average size of about 200 (Figure 6D-E) or 100 mm^3^ (Figure 7A-B) the mice were treated 3 times a week with osimertinib (5 mg/kg for experiments with PC9 cells; 10 mg/kg for experiments with YUX-1024 cells) and sorafenib (60 mg/kg), alone or in combination, using a water/EtOH/Kolliphor (Sigma-Aldrich) solution. For the experiment shown in Figure 6B, the mice were initially treated 5 times a week for 12 days, before switching to a 3-times-a-week regimen for the rest of the experiment. When the size of at least one of the tumors reached the arbitrary volume of 800 mm^3^, the mice were sacrificed and the tumors were dissected for gDNA extraction.

For iDISCO 3D-analysis, PC9 cells containing the EGFR-C797S, PIK3CA-E545K or KRAS-G12D CRISPR-barcodes were selected with osimertinib (0.1 µM) and transduced with GFP, mCherry or b-galactosidase lentiviruses, respectively. The cells were selected with puromycin, and GFP and mCherry containing cells were sorted by FACS. The three cell populations were mixed with parental PC9 cells in a 1:1000 proportion, to form a mass population of osimertinib-sensitive (unlabeled) and osimertinib-resistant (labeled) cells. YUX-1024 cells containing the BRAF-V600E CRISPR-barcode and selected with osimertinib (0.1 µM) were transduced with a GFP lentivirus, puromycin selected and FACS sorted. The cells were then mixed with parental YUX-1024 (1:200 ratio). PC9 and YUX-1024 cells were mixed with Matrigel and subcutaneously inoculated in the flanks of female SCID mice. When the tumors reached an average size of 200 mm^3^, the mice were randomized and treated 3 times a week with osimertinib (5 mg/kg for the PC9 experiment, 10 mg/kg for the YUX-1024 experiment) alone or in combination with sorafenib (60 mg/kg) by gavage. Control mice were sacrificed after randomization. The other mice were sacrificed after 4 weeks of treatment and the tumors were fixed overnight at 4°C in 4% paraformaldehyde.

BEM-5 cells were generated by introducing through CRISPR-barcoding a subpopulation of EGFR-C799S (corresponding to EGFR-C797S in the human receptor) cells in BALB-3T3-EGFR-L860R-clone M5. BEM-5 cells were bilaterally injected in the flanks of female BALB/c mice (2 × 10^6^ cells per site). When the tumors reached an average size of 50 mm^3^, the mice were treated 4 times a week with osimertinib (20 mg/kg) or 3 times a week with sorafenib (60 mg/kg), alone or in combination with osimertinib.

Animal experiments were approved by Mount Sinai’s Animal Care and Use Committee (IACUC) or the French Regional Ethics Committees and the Ministry of Education, Research and Innovation (project n°19253-201903191709966 v1 and authorization n° 11752 – 16/10/18), and they were performed in accordance with the European Committee Council Directive (2010-63-EU).

### iDISCO 3D-imaging

Sample processing, immunostaining and imaging were described previously (Belle et al., 2014; Belle et al., 2017). Briefly, the tumors were dehydrated with increasing concentrations of MetOH (20, 40, 60, 80 and 100%), followed by incubation in a solution of 2/3 dichloromethane (DCM; Sigma-Aldrich), 1/3 MetOH. The tumors were then bleached overnight using a fresh 5% solution of H_2_O_2_ in MetOH at 4°C, followed by rehydration with decreasing concentrations of MetOH (100, 80, 60, 40 and 20%). The tumors were incubated for 2 days with a permeabilizing solution containing PBS1, Triton-X 100, glycine and 10% DMSO, then incubated for two additional days with a blocking solution containing 6% Donkey Serum, Triton-X 100 and 10% DMSO. The anti-GFP, anti-RFP/mCherry and anti-beta-galactosidase antibodies (Table S7) were incubated in PTwH/5%DMSO/3% Donkey Serum for 1 week at 37°C. After 5 washes, the tumors were incubated with the secondary antibodies (Table S7) in PTwH/3% Donkey Serum for 1 week at 37°C. The tumors were washed 5 times and then dehydrated using increasing concentrations of MetOH, followed by a delipidation using DCM solutions and overnight clearing in dibenzylether (DBE; Sigma-Aldrich).

Cleared samples were imaged with an Ultramicroscope II (LaVision BioTec) using the ImspectorPro software (LaVision BioTec). The light sheet was generated by a Coherent Sapphire Laser (LaVision BioTec) at wavelengths of 488, 561and 640 nm and six cylindrical lenses. A binocular stereomicroscope (MXV10, Olympus) with a 2x objective (MVPLAPO, Olympus) was used at different magnifications (0.63X and 0.8x). The samples were placed in an imaging reservoir made of 100% quartz (LaVision BioTec), filled with DBE and illuminated from the side by the laser light. Images were acquired with a PCO Edge SCMOS CCD Camera (2560 x 2160 Pixel size, LaVision BioTec). The step size in Z-orientation between each image was fixed at 4 or 6 µm for 0.63x and 0.8x magnifications. Images were processed and analyzed using the Imaris software (Bitplane).

### Immunohistochemical analysis of BALB/c tumors

Immunohistochemistry was performed on formalin fixed and deparaffinized tissue sections. Sections were heated at 95 °C for 20 min in 10 mM citrate buffer (pH 6) and/or Tris EDTA (pH 9) for antigen retrieval, treated with peroxidase blocking reagent (Dako) and incubated with primary antibodies at the concentration indicated in Table S7. The sections were incubated with secondary antibodies coupled to peroxidase and revealed with diaminobenzidine (Dako). The tissue sections were counterstained with hematoxylin and images were acquired on an Axioscope 7 microscope (Zeiss).

### Statistical analysis

Statistical analysis was performed with the GraphPad Prism software using Mann-Whitney’s test.

## ACKNOWLEDGMENTS

We thank M. Thomas, I. Lihrmann, T. Lecroq, L. Mouchard for helpful discussion. We thank P. Jänne (Dana-Farber Cancer Institute), J. Minna (UT Southwestern Medical Center) and R. Whitehead (Vanderbilt University) for sharing cell lines. Some experiments and analyses, including part of the qPCR for the initial screen, amplicon quality assessment by Bioanalyzer, 3D-imaging analysis using the Imaris software and acquisition of immunostained sections, were performed at the Cell Imaging Platform of Normandie (PRIMACEN). We thank D. Cartier (NorDiC), G. Riou (IRIB Flow Cytometry Facility), J. Maucotel (SRB, UNIROUEN), M. Di Giovanni, M. Bénard and A. Lebon (PRIMACEN) for technical assistance.

This work was supported by the Institut National de la Santé et de la Recherche Médicale (INSERM), the Université de Rouen Normandie, the Institut National du Cancer (PLBIO 2017-159), the Agence Nationale de la Recherche (METROPOLIS), the Ligue contre le Cancer de Haute-Normandie, the Fondation ARC pour la Recherche sur le Cancer (PJA20161205119), the Conseil Régional de Normandie and the FEDER program of the European Union. L.B. was supported by a doctoral fellowship from the Ligue contre le Cancer de Normandie and the Normandie Region. A.G. was supported by a doctoral fellowship from the Normandie Region. H.C. was supported by doctoral fellowships from the Nabatieh Municipal Council (Lebanon) and the Fondation pour la Recherche Médicale. B.C.C. was supported by Basic Science Research Program through the NRF funded by the Korean Ministry of Science and ICT (2016R1A2B3016282).

## AUTHOR CONTRIBUTIONS

L.B., D.A., J.L., MdM.B.-R., A.G., H.C., Z.K., A.A., S.Y., D.G., J.D., P.J., A.V. and L.G. performed the experiments. L.B., D.A., J.L., MdM.B.-R., A.G., H.C., Z.K., A.A., C.D., D.T., S.A.A., A.B.C., Y.A. and L.G. designed and/or analyzed experiments. J.-R. L. and C.Cheng analyzed the EGFR-score. C.B. participated in the analysis of highly complex CRISPR barcode. S.A. and A.B. provided CRC cell lines and participated in experiment design and analysis. C.Coulouarn performed and analyzed gene array experiments. B.C.C. provided the PDC lines and participated in experiment design and analysis. L.G., Y.A. and A.B.C. acquired funding. S.A.A, S.A., D.T., Z.K. and Y.A. edited the manuscript. L.G. conceived and supervised the study, and wrote the manuscript.

## DECLARATION OF INTERESTS

Z.K. reports financial support from DeuterOncology NV outside the submitted work. B.C.C. reports stock ownership with TheraCanVac Inc, Gencurix Inc, Bridgebio therapeutics, KANAPH Therapeutic Inc, Cyrus Therapeutics, Interpark Bio Convergence Corp and J INTS BIO; reports participating in an advisory role for KANAPH Therapeutic Inc, Brigebio Therapeutics, Cyrus Therapeutics, Guardant Health and Oscotec; has received consulting fees from Novartis, Abion, BeiGene, AstraZeneca, Boehringer-Ingelheim, Roche, Bristol Myers Squibb, ONO, Yuhan, Pfizer, Eli Lilly, Janssen, Takeda, MSD, Janssen, Medpacto, and Blueprint medicines; has received grants or funds from Novartis, Bayer, AstraZeneca, MOGAM Institute, Dong-A ST, Champions Oncology, Janssen, Yuhan, ONO, Dizal Pharma, MSD, Abbvie, Medpacto, GI Innovation, Eli Lilly, Blueprint medicines, and Interpark Bio Convergence Corp; has received royalties from Campions Oncology, Crown Bioscience and Imagen; and is the founder of DAAN Biotherapeutics. A.B.C. has received honorarium for advisory positions, board memberships, lectures or non-financial support from the following sources: Astra-Zeneca, Roche, MSD, Pfizer, Novartis, Takeda, Janssen, Abbvie, Amgen. L.G. is inventor of a patent on DNA barcoding issued to Inserm and University of Rouen.

---

**Table S4.**
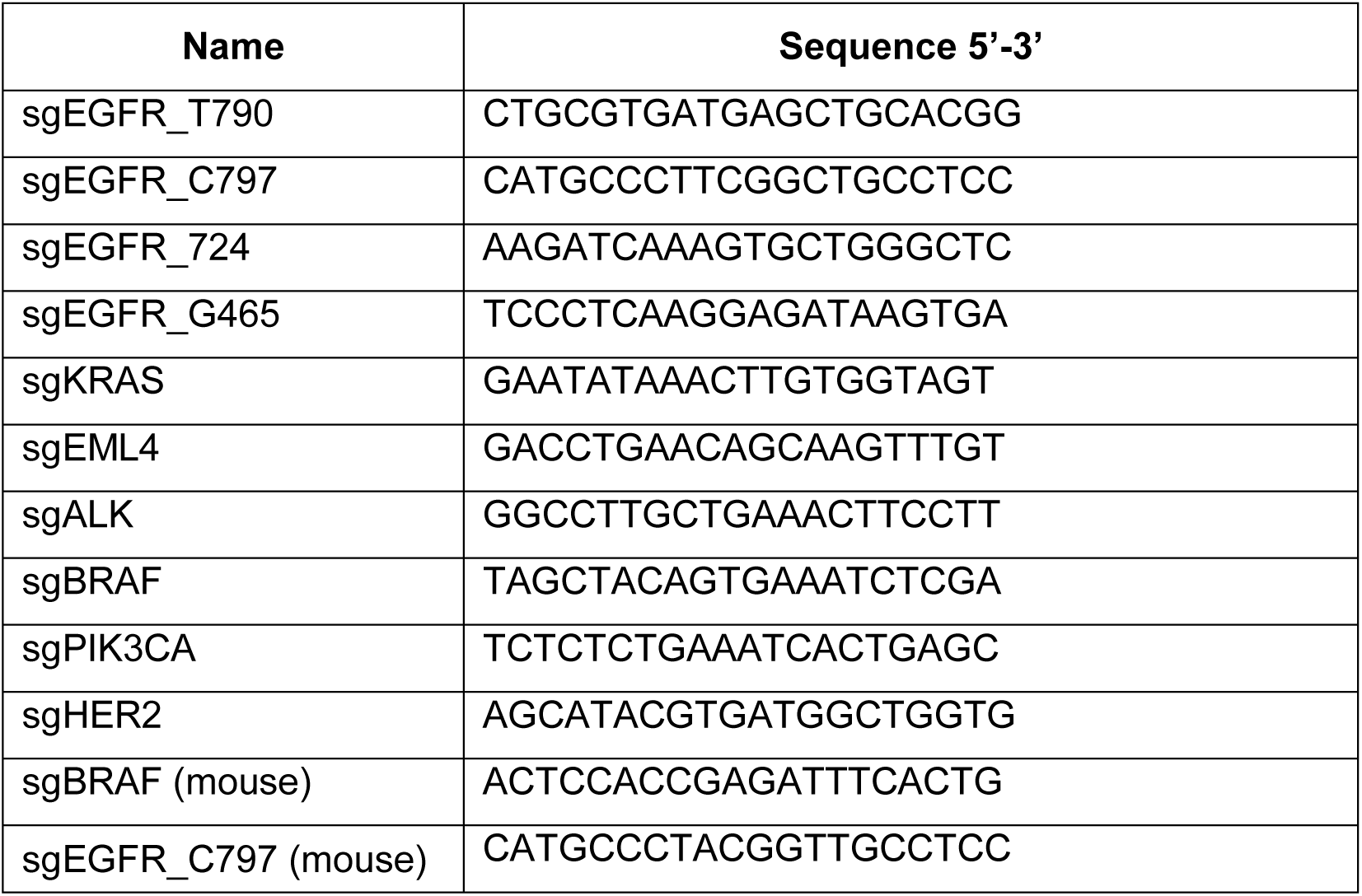
List of sgRNA target sequences used for CRISPR-barcoding.

**Table S5.**
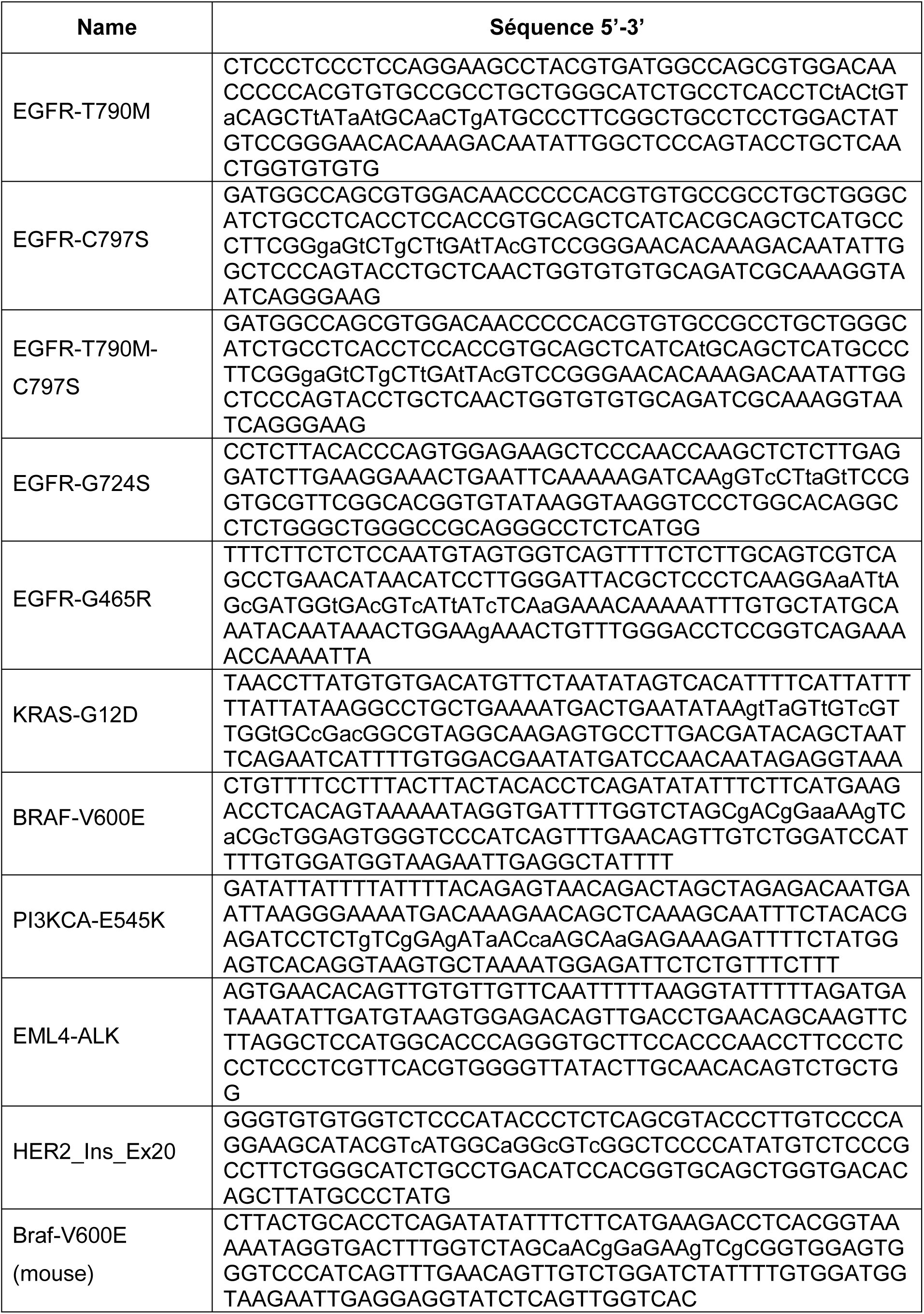
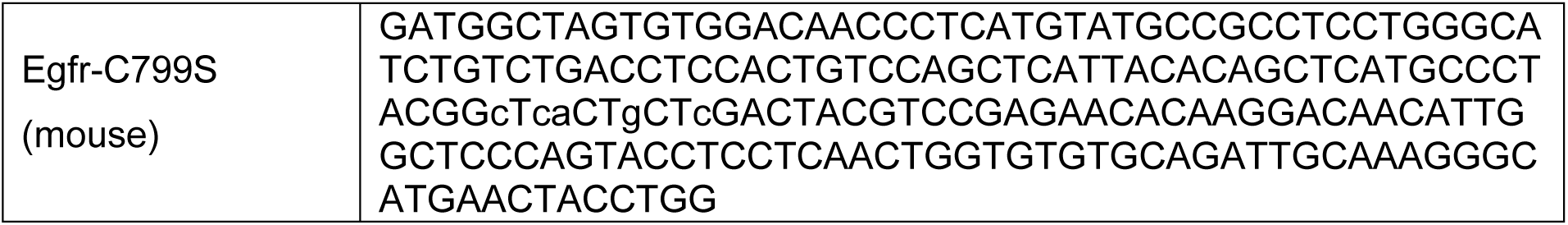
List of ssODNs used for CRISPR-barcoding. Mutations compared to the endogenous sequence are indicated in lowercase letters.

**Table S6.**
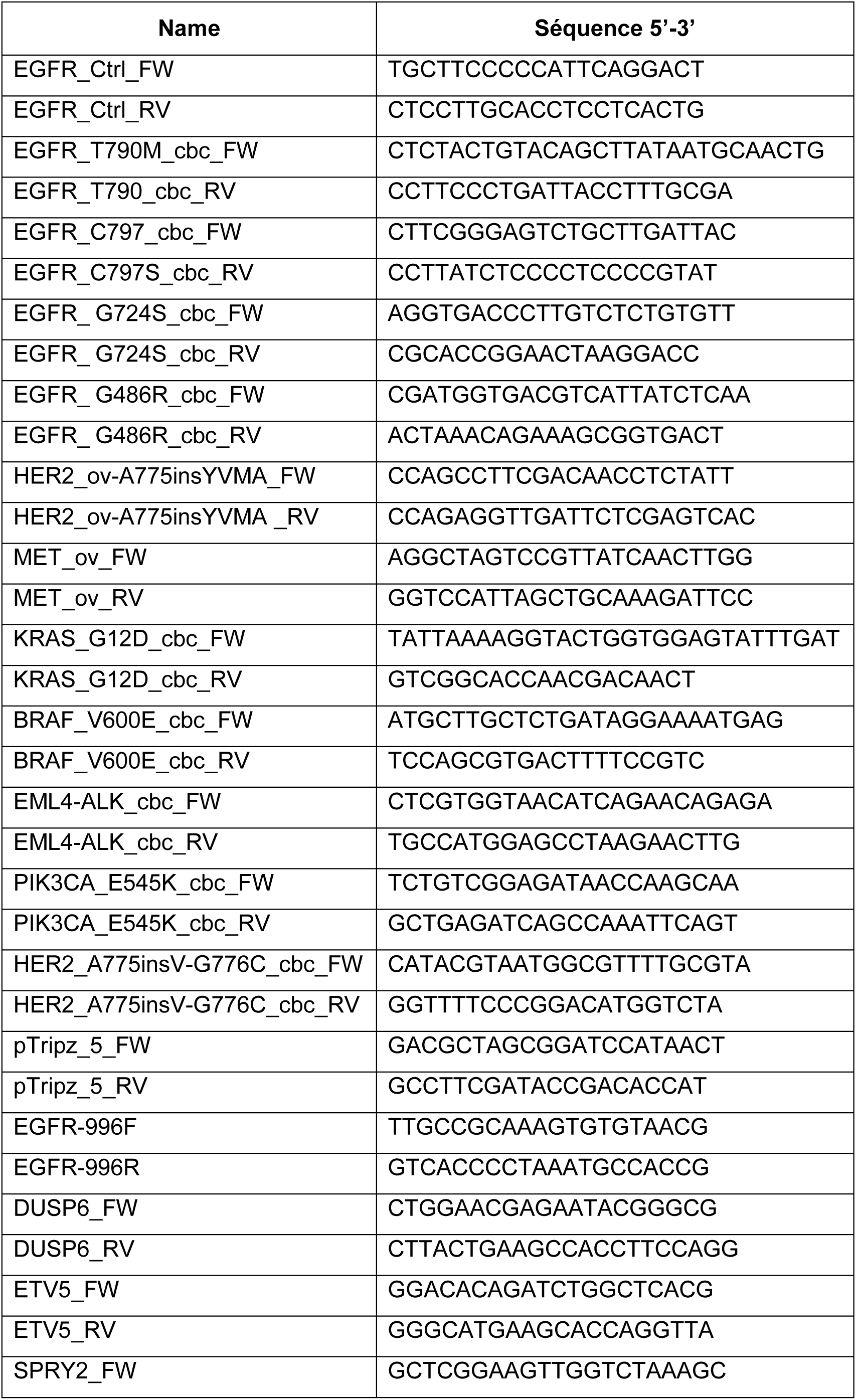
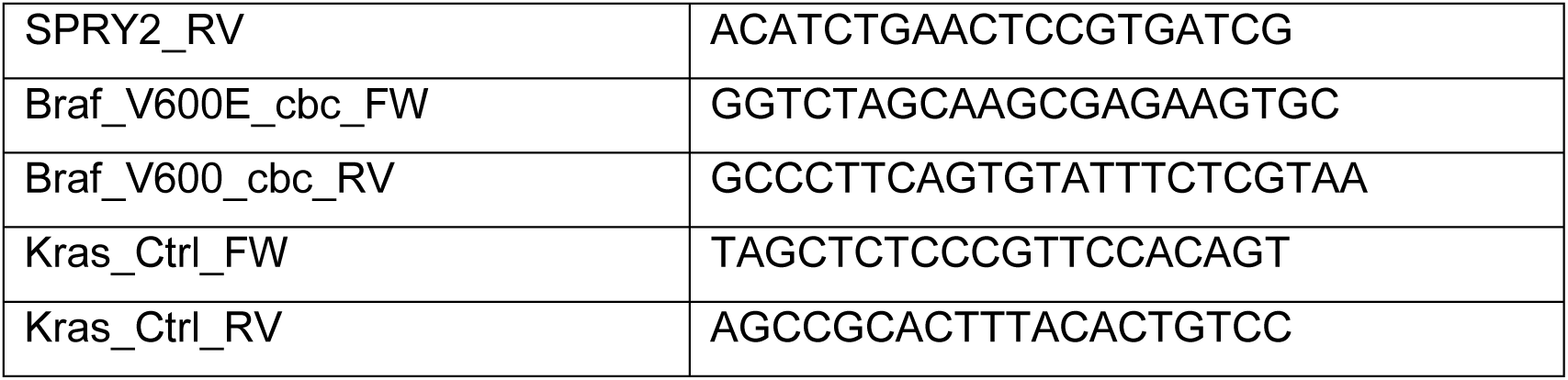
List of primers used for qPCR.

**Table S7.** List of the antibodies used.

| **Antibody** | **Reference** | **Dilution** |
| --- | --- | --- |
| Akt (pan) (40D4) - Mouse mAb | #2920, Cell Signaling | 1:1,000 |
| c-Myc Clone 9E10, Mouse mAb | 11667149001, Roche | 1:500 |
| EGF Receptor - Rabbit | #2232, Cell Signaling | 1:1,000 |
| eIF4E (C46H6) - Rabbit mAb | #2067, Cell Signaling | 1:1,000 |
| FLAG M2, Clone M2 - Mouse mAb | F1804, Sigma | 1:500 |
| GAPDH - Mouse mAb | G8795, Sigma | 1:5,000 |
| HER2/ErbB2 (D8F12) - Rabbit mAb | #4290, Cell Signaling | 1:1,000 |
| Mcl-1 (D35A5) - Rabbit mAb | #5453, Cell Signaling | 1:1,000 |
| Met (D1C2) - Rabbit mAb | #8198, Cell Signaling | 1:1,000 |
| MNK2 - Rabbit | ab84345, Abcam | 1: 1,000 |
| p44/42 MAPK (Erk1/2) (L34F12) Mouse mAb | #4696, Cell Signaling | 1:1,000 |
| Phospho-p44/42 MAPK (Erk1/2) (Thr202/Tyr204) (D13.14.4E) Rabbit mAb | #4370, Cell Signaling | 1:1,000 |
| Phospho-Akt (Ser473) (D9E) - Rabbit mAb | #4060, Cell Signaling | 1:1,000 |
| Phospho-EGF Receptor (Tyr1068) (D7A5) - Rabbit mAb | #3777, Cell Signaling | 1:1,000 |
| Phospho-eIF4E (Ser209) - Rabbit | #9139, Cell Signaling | 1:1,000 |
| Phospho-S6 Ribosomal Protein (Ser240/244) | #2215s, Cell Signaling | 1:1,000 |
| Phospho-Stat3 (Tyr705) (D3A7) - Rabbit mAb | #9145, Cell Signaling | 1:1,000 |
| Phospho-Stat3 (Tyr705) – Rabbit | #9131, Cell Signaling | 1:1,000 |
| Stat3 (124H6) - Mouse mAb | #9139, Cell Signaling | 1:1,000 |
| Slug – Rabbit mAb | #9585s, Cell Signaling | 1:1,000 |
| Tubulin - Mouse mAb | T5168, Sigma | 1:5,000 |
| Vimentin - Rabbit | #3932, Cell Signaling | 1:1,000 |
| β-galactosidase - Goat polyclonal | 0856028, MP Biomedicals | 1:1,000 |
| GFP - Chicken polyclonal | GFP-1010, Aves Labs | 1:2,000 |
| RFP - Rabbit polyclonal | 600-401-379, Rockland | 1:500 |
| Anti-Goat Alexa Fluor 488 | 705-545-147, Jackson ImmunoResearch | 1:800 |
| Anti-Chicken Cyanine 3 | 703-165-155, Jackson ImmunoResearch | 1:800 |
| Anti-Rabbit Alexa Fluor 647 | 711-605-152, Jackson ImmunoResearch | 1:800 |
| Anti-Chicken Alexa Fluor 647 | 703-605-155, Jackson ImmunoResearch | 1:800 |
| Anti-CD8 | #98941, Cell Signaling | 1:300 |
| Anti-CD3 | MCA1477, Bio-Rad | 1:100 |
| Anti-CD68 | #97778, Cell Signaling | 1:400 |
| Anti-CD86 | #19589, Cell Signaling | 1:400 |

