## Supplementary material for "Prolonging lung cancer response to EGFR inhibition by targeting the selective advantage of resistant cells": Suppl. Table1

**Table S1. Small molecule screen in CRISPR-barcoded PC9 cells treated in combination with gefitinib (Gef).**

| | | | EGFR T790M | | | | EML4-ALK | | | | KRAS G12D | | | |
|---|---|---|---|---|---|---|---|---|---|---|---|---|---|---|
| | | | EXP1 | | EXP2 | | EXP1 | | EXP2 | | EXP1 | | EXP2 | |
| | **Concentrations** | **Target** | **mean** | *sem* | **mean** | *sem* | **mean** | *sem* | **mean** | *sem* | **mean** | *sem* | **mean** | *sem* |
| Ctrl | | | **1** | *0,15* | **1** | *0,25* | **1,03** | *0,16* | **1** | *0,04* | **1,01** | *0,12* | **1** | *0,05* |
| Gef | 1 uM | EGFR | **4,21** | *1,33* | **4,77** | *0,71* | **2,71** | *0,09* | **4,15** | *0,17* | **3,24** | *0,21* | **3,42** | *0,13* |
| Gef + TAE684 | 0,5 uM | ALK | **4,97** | *1,01* | **3,47** | *1,51* | **1,13** | *0,06* | **1,36** | *0,06* | **4,76** | *0,22* | **7,99** | *2,83* |
| Gef + WZ42002 | 0,5 uM | EGFR (3rd gen) | **0,7** | *0,18* | **0,8** | *0,06* | **2,44** | *0,15* | **3,12** | *0,24* | **3,31** | *0,24* | **4,36** | *1,56* |
| Gef + KU55933 | 5 uM | ATM | **5,05** | *0,46* | **2,56** | *0,48* | **3,26** | *0,23* | **4,56** | *0,56* | **4,07** | *0,43* | **8,84** | *2,39* |
| Gef + IWP2 | 5 uM | PORCN (Wnt) | **3,27** | *1,05* | **4,32** | *1,25* | **2,82** | *0,37* | **3,89** | *0,27* | **3,58** | *0,5* | **4,73** | *0,93* |
| Gef + AZD5363 | 1 uM | AKT | **2,61** | *0,24* | **1,68** | *0,63* | **2,38** | *0,12* | **2,03** | *0,1* | **2,73** | *0,35* | **2,14** | *0,11* |
| Gef + Sunitinib | 1 uM | Multikinase | **3,32** | *0,6* | **3,98** | *0,75* | **1,68** | *0,27* | **1,97** | *0,36* | **2,99** | *0,26* | **4,05** | *0,74* |
| Gef + Sorafenib | 1 uM | Multikinase | **1,15** | *0,03* | **1,15** | *0,14* | **1,86** | *0,11* | **3,16** | *0,15* | **2,21** | *0,04* | **1,87** | *0,19* |
| Gef + Compound C | 1 uM | AMPK | **2,68** | *0,58* | **3,22** | *0,66* | **2,62** | *0,23* | **4,29** | *0,22* | **4,22** | *0,52* | **3,58** | *0,27* |
| Gef + U0126 | 1 uM | MEK1/2 | **3,12** | *0,51* | **5** | *2* | **3,59** | *0,17* | **4,19** | *0,2* | **4,1** | *0,11* | **6,83** | *1,02* |
| Gef + STO-609 | 1 uM | CaMK | **4,64** | *1,06* | **2,79** | *0,38* | **3,04** | *0,27* | **4,19** | *0,11* | **3,71** | *0,63* | **4,71** | *0,67* |
| Gef + PS1145 | 1 uM | IKK (Nf-κB) | **3,53** | *0,84* | **2,57** | *0,75* | **2,92** | *0,22* | **5,23** | *1,25* | **3,27** | *0,18* | **5,79** | *2,16* |
| Gef + H89 | 1 uM | PKA | **2,48** | *0,15* | **2,43** | *0,56* | **2,05** | *0,4* | **3,67** | *0,1* | **3,32** | *0,29* | **5,04** | *0,98* |
| Gef + Chelerythrin | 1 uM | PKC | **2,34** | *0,38* | **1,87** | *0,09* | **2,46** | *0,09* | **3,85** | *0,13* | **3,52** | *0,15* | **2,65** | *0,32* |
| Gef + Doxorubicin | 50 nM | DNA topoisomerase II | **1,09** | *0,27* | **4,52** | *1,08* | **1,3** | *0,07* | **1,79** | *0,27* | **1,76** | *0,26* | **2,25** | *0,44* |
| Gef + Verteporfin | 2 uM | YAP | | | **5,47** | *1,08* | | | **4,06** | *0,47* | | | **5,59** | *0,85* |
