## Supplementary material for "Prolonging lung cancer response to EGFR inhibition by targeting the selective advantage of resistant cells": Suppl. Figures

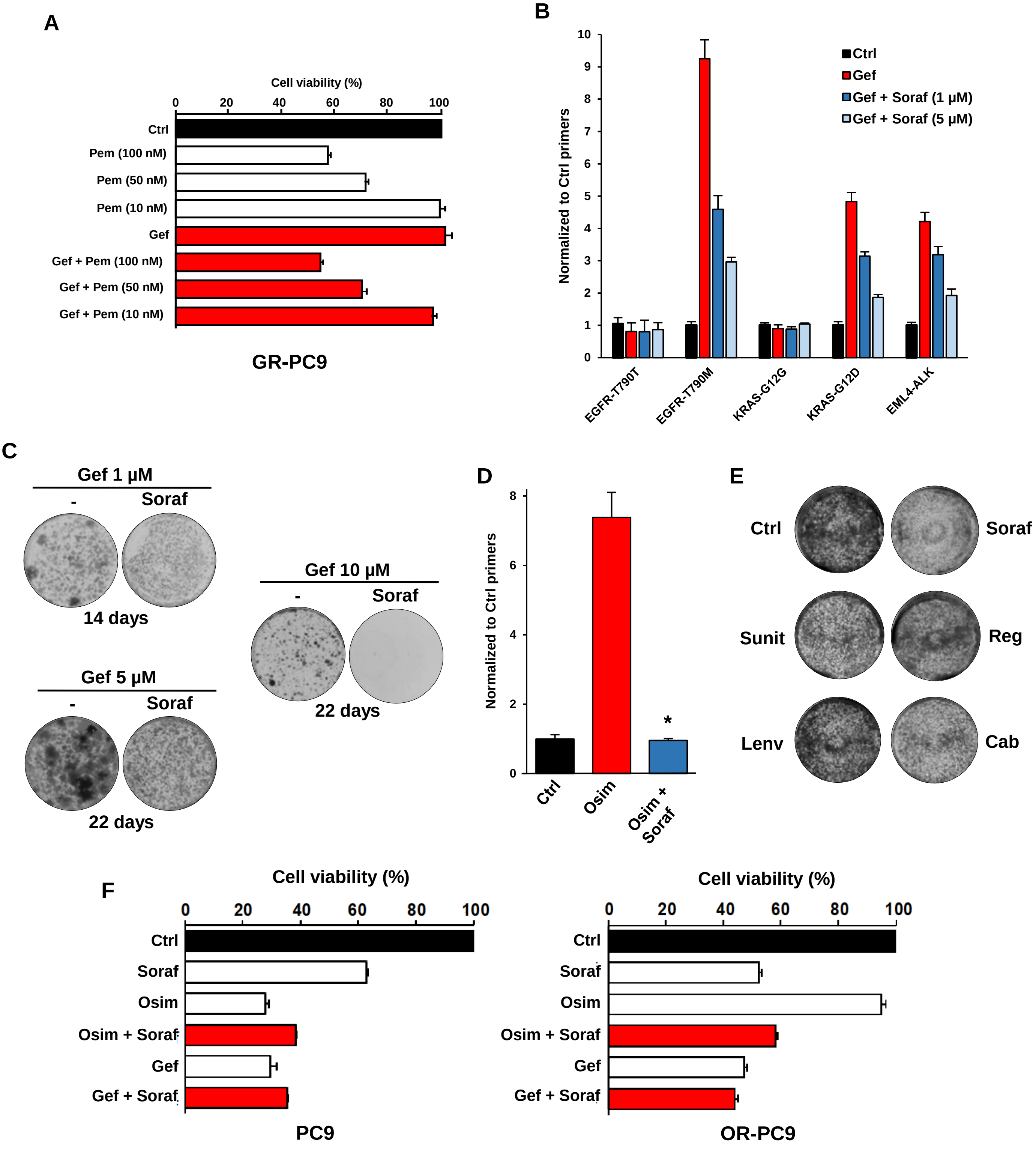


**Figure S1. Effects of pemetrexed and sorafenib on the emergence of resistant subpopulations of NSCLC cells induced by EGFR-TKIs. A**, Cell viability assay of gefitinib-resistant (EGFR-T790M) PC9 cells grown for 5 days in the presence or the absence of gefitinib (1 µM) or pemetrexed (100, 50 or 10 nM). The fraction of viable cells was measured by CellTiter-Glo and normalized to the DMSO-treated control. **B**, The indicated CRISPR-barcodes were introduced in PC9 cells, and the cells were treated with or without gefitinib (1 µM), alone or in combination with sorafenib (1 or 5 µM), for 5 days. The proportion of the barcodes was measured by qPCR from genomic DNA and normalized using EGFR_Ctrl primers. **C**, Representative images of colony forming assays of the same population of CRISPR-barcoded PC9 cells treated with gefitinib (1, 5 or 10 µM) alone or in combination with sorafenib (5 µM) for 14 or 22 days. **D**, CRISPR-barcoding was used to introduce the EGFR-C797S barcode in a population of PC9 cells containing the EGFR-T790M mutation. The cells were then treated with osimertinib (1 µM) or in combination with sorafenib (5 µM) for 9 days, and the proportion of the EGFR-C797S barcode was assessed by qPCR. The mean ±SEM (n=4) of one representative of three experiments is represented. **E**, Colony forming assay of PC9 cells treated for 5 days with sorafenib (5 µM), sunitinib (1 µM), regorafenib (2 µM), lenvatinib (1 µM) or cabozantinib (5 µM). **F**, Cell viability assay of parental or osimertinib-resistant PC9 cells grown for 5 days in the presence or the absence of gefitinib (1 µM) or osimertinib (0,1 µM) alone or in combination with sorafenib (5 µM).

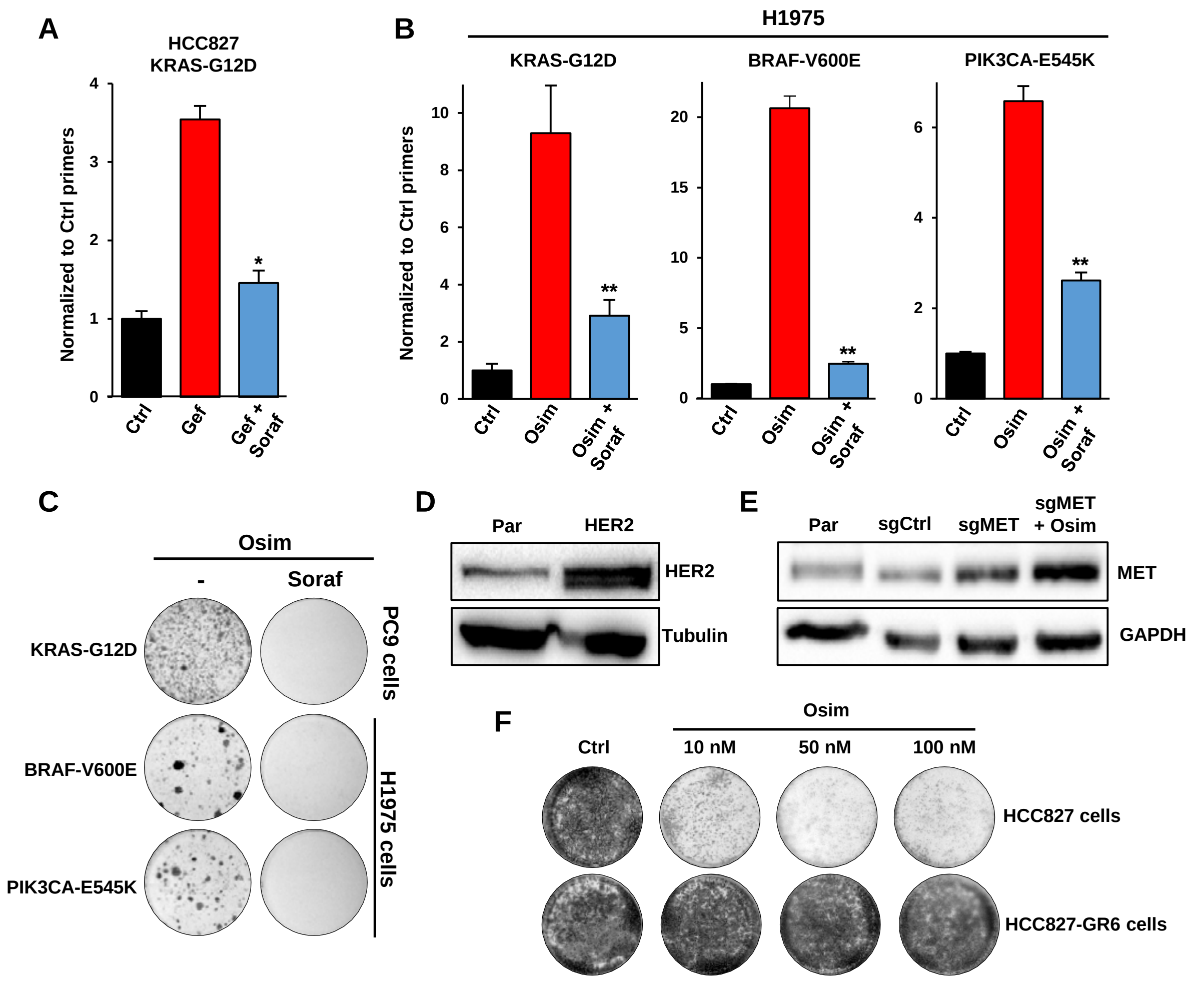


**Figure S2. Effects of Sorafenib on different mechanisms of resistance to EGFR-TKIs. A**, Effects of gefitinib (1 µM), with or without sorafenib (5 µM), on the proportion of HCC827 cells containing the CRISPR-barcode KRAS-G12D after 5 days of treatment. The barcode levels were assessed by qPCR and normalized using EGFR_Ctrl primers. Mean (±SEM; n=4) of one representative of three experiments. **B**, Effects of osimertinib (0,1 µM) with or without sorafenib (5 µM) on the proportion of KRAS-G12D (15 days), BRAF-V600E (7 days) or PIK3CA-E545K (15 days) CRISPR-barcodes in H1975 cells. Mean (±SEM; n=4) of individual experiments repeated at least three times. **C**, Representative images of colony forming assays of PC9 and H1975 cells containing the indicated CRISPR-barcodes and treated with osimertinib alone (0,1 µM) or in combination with sorafenib (5 µM) for one month. **D**, PC9 cells were transduced (HER2) or not (Par) with a HER2 lentivirus and immunoblot was performed using the indicated antibodies. **E**, The levels of MET receptor were assessed by immunoblot in parental (Par) PC9 and in cells containing the dCas9 activator system with a control (sgCtrl) or a MET-specific (sgMET) sgRNA. A population of sgMET cells was selected for two weeks with osimertinib (0,1 µM; sgMET+Osim) before the lysis. **F**, HCC827 and HCC827-GR6 cells were treated for 7 or 5 days, respectively, with the indicated concentrations of osimertinib, followed by fixation and Crystal violet staining.

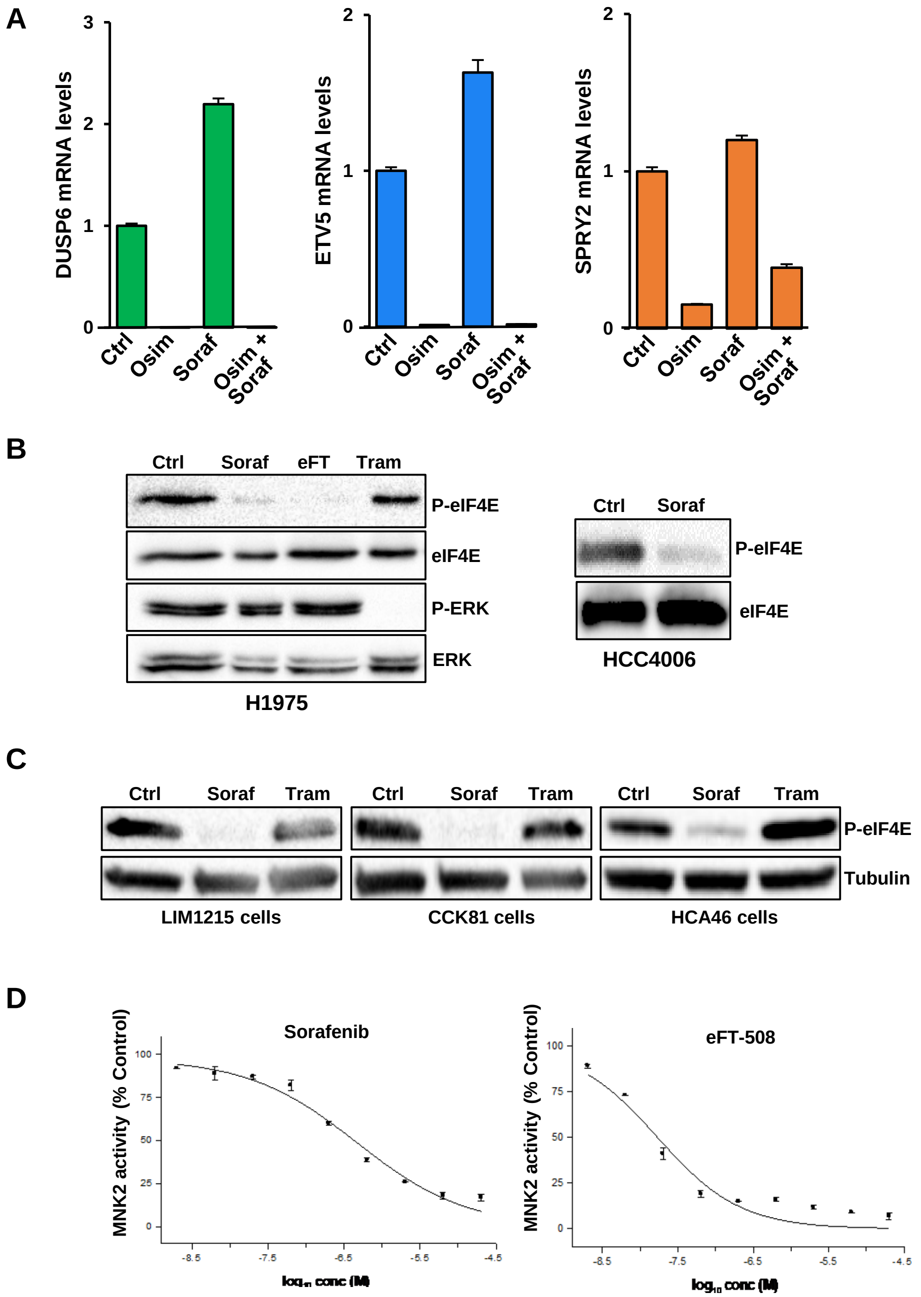


E

| Assay | Compound | IC50 |
|---|---|---|
| MNK1 | Sorafenib | 6.0E-06 M |
| | eFT-508 | 2.4E-09 M |
| MNK2 | Sorafenib | 4.62E-07M |
| | eFT-508 | 1.7E-08M |

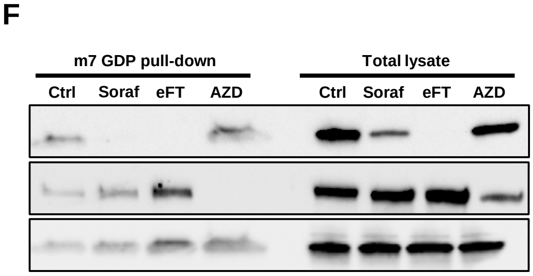


**Figure S3. Sorafenib blocks phosphorylation of eIF4E by a direct inhibition of MNK activity without affecting MAPKs in NSCLC cells. A**, PC9 cells were treated with osimertinib (1 µM) and sorafenib (5 µM), alone or in combination, for 2 days and the expression of DUSP6, SPRY2 and ETV5 was assessed by qPCR. **B**, H1975 and HCC4006 NSCLC cells were treated for 2h with or without sorafenib (5 µM), eFT-508 (1 µM; eFT, MNK inhibitor) or trametinib (50 nM; Tram, MEK inhibitor), followed by immunoblot using the indicated antibodies. **C**, LIM1215, HCA46 and CCK81 CRC cells were treated for 2h in the presence or the absence of sorafenib (3 µM) or trametinib (25 nM), followed by immunoblot using anti-phospho-eIF4E or anti-tubulin antibodies. **D**, Effects of sorafenib and eFT-508 on the *in vitro* catalytic activity of MNK2, measured using as a substrate the myelin basic protein. **E**, Table reporting the *in vitro* IC50 of sorafenib and eFT-508 for MNK1 and MNK2, measured using as a substrate the myelin basic protein or a peptide derived from the human cAMP Response Element Binding protein. **F**, Cap pull-down assay in PC9 cells treated for 6h in the presence or the absence of sorafenib (5 µM), eFT-508 (1 µM) or AZD8055 (1 µM; AZD, mTOR inhibitor). Cell lysate was pulled-down using m7 GDP beads, and immunoblot was performed using the indicated antibodies.

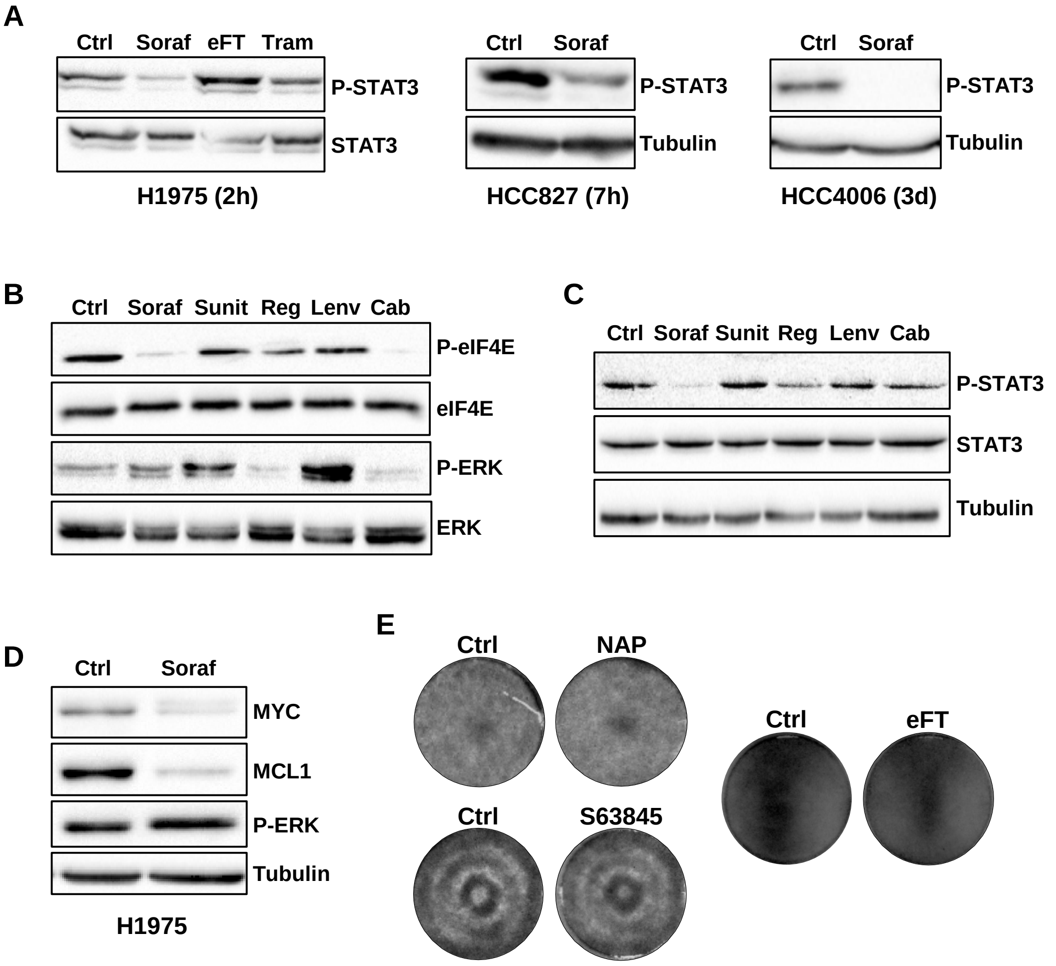


**Figure S4. Sorafenib inhibits phosphorylation of STAT3 and expression of MYC and MCL1 in NSCLC cells. A**, H1975, HCC827 and HCC4006 cells were treated for the indicated time points with sorafenib (5 µM), eFT-508 (1 µM) or trametinib (50 nM), followed by immunoblot using anti-phospho-STAT3, anti-STAT3 or anti-tubulin antibodies. **B**, **C**, PC9 cells were treated for 2h (**B**) or 6h (**C**) with sorafenib (5 µM), sunitinib (1 µM), regorafenib (2 µM), lenvatinib (1 µM) or cabozantinib (5 µM) and immunoblot was performed using the indicated antibodies. **D**, H1975 cells were treated for 3 days in the presence or the absence of sorafenib (5 µM), followed by immunoblot using the indicated antibodies. **E**, Representative images of colony forming assays of PC9 cells treated for 6 days with napabucasin (0,5 µM; NAP, STAT3 inhibitor), S63845 (0,1 µM; MCL1 inhibitor) or eFT-508 (1 µM).

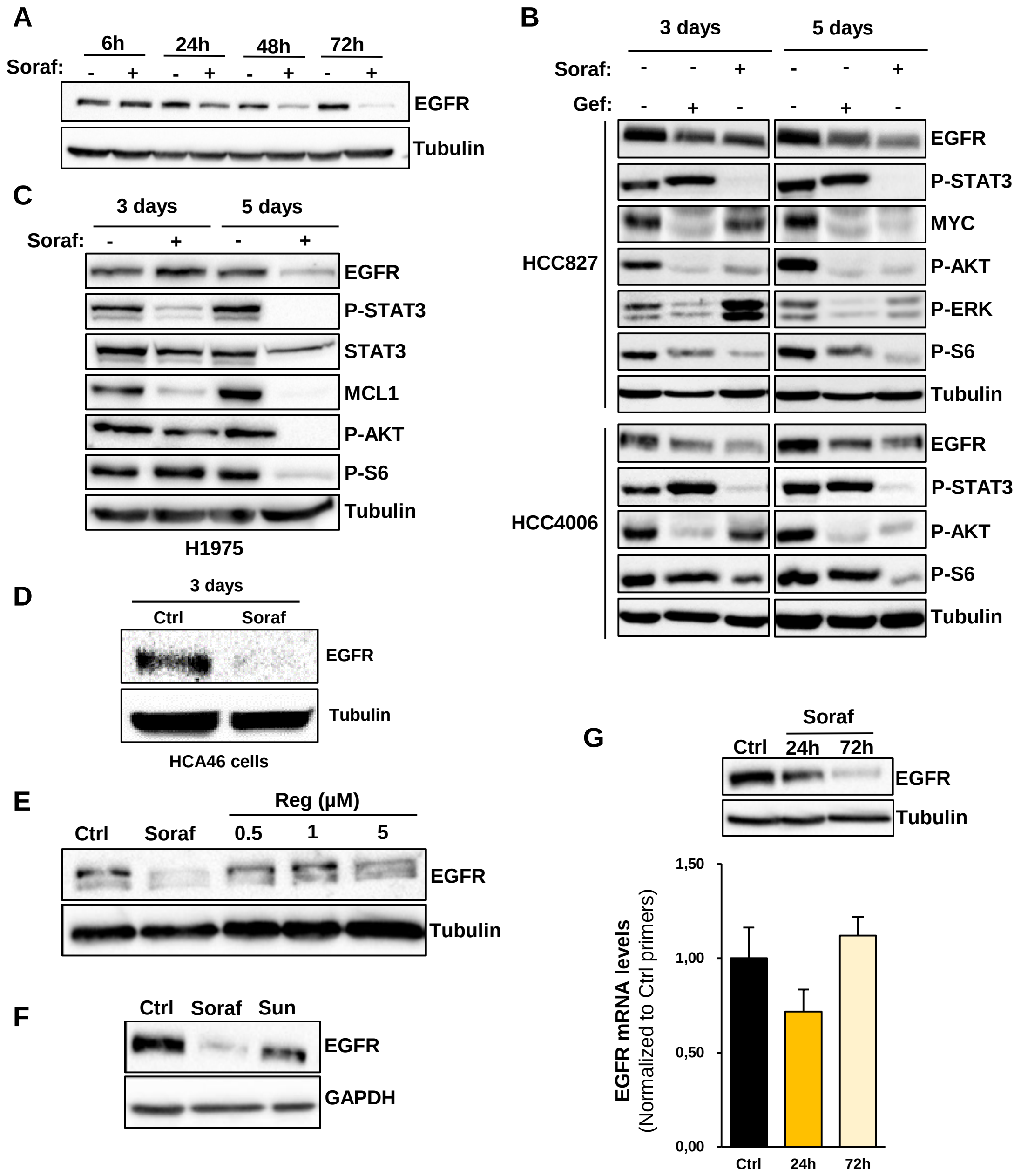


**Figure S5. EGFR down-regulation by Sorafenib in NSCLC cells. A,** PC9 cells were treated for the indicated time points with sorafenib (5 µM) and immunoblot was performed with anti-EGFR and anti-tubulin antibodies. **B**, HCC827 and HCC4006 NSCLC cells were treated for 3 or 5 days in the presence or the absence of sorafenib (5 µM) or gefitinib (1 µM), followed by immunoblot using the indicated antibodies. **C**, H1975 NSCLC cells were treated for 3 and 5 days with or without sorafenib (5 µM), followed by immunoblot using the indicated antibodies. **D**, HCA46 CRC cells were treated with sorafenib (5 µM) for 3 days, followed by immunoblot using anti-EGFR and anti-tubulin antibodies. **E**, PC9 cells were treated for 4 days with or without sorafenib (5 µM) or regorafenib, followed by immunoblot (lower panel). Colony forming assay of PC9 cells treated with increasing concentrations of regorafenib (upper panel). **F,** PC9 cells were treated for 3 days with sorafenib (5 µM) or sunitinib (1 µM), followed by immunoblot with anti-EGFR and anti-GAPDH antibodies. **G**, PC9 cells were treated for 24 and 72h with sorafenib (5 µM) and the levels of EGFR mRNA and protein were assessed by immunoblot (upper panel) or RT-qPCR (lower panel).

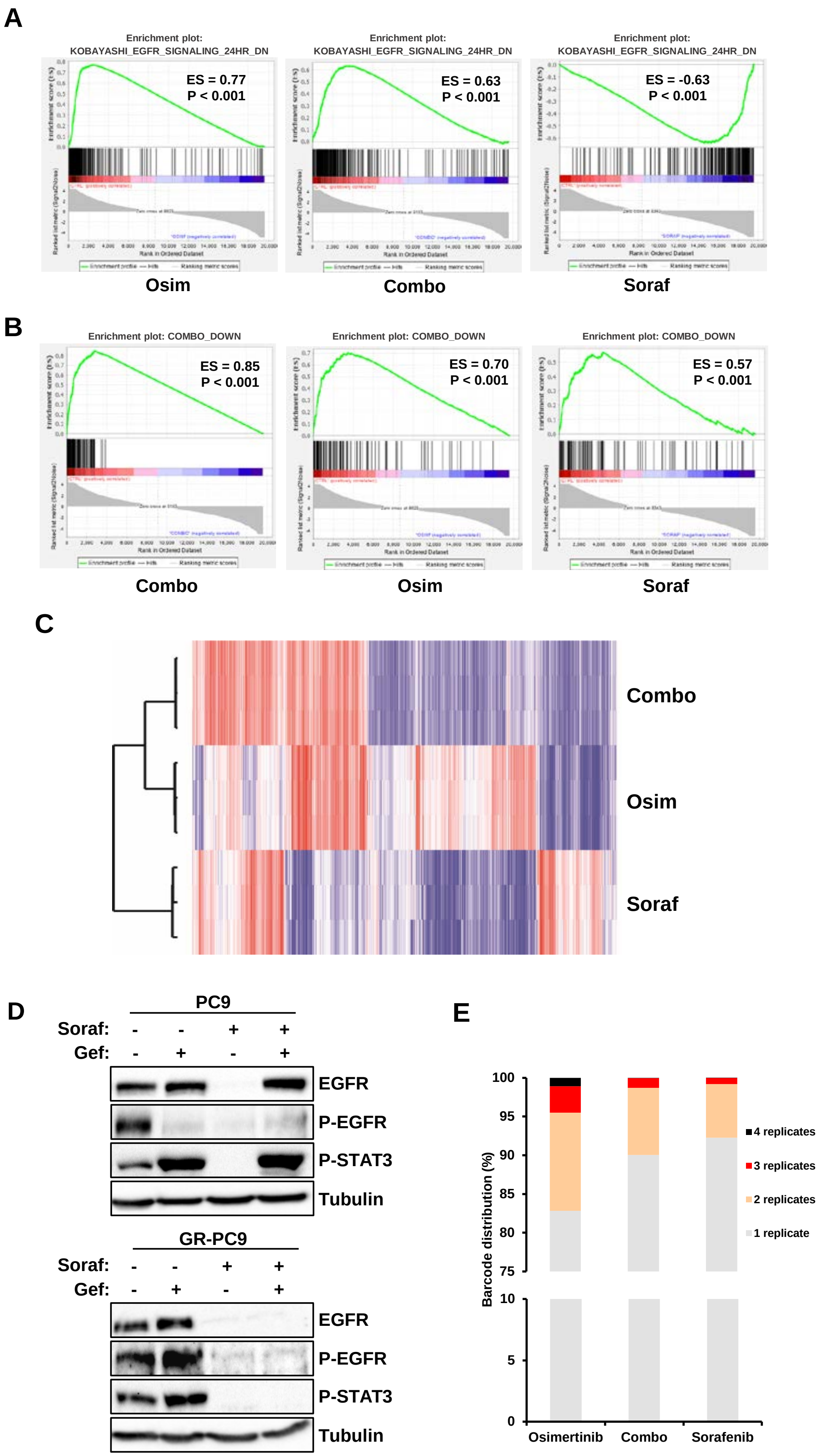


**Figure S6. Effects of sorafenib on the clonal evolution of NSCLC cells induced by osimertinib. A**, Gene set enrichment analysis (GSEA) of EGFR-TKI down-regulated genes in NSCLC cells (KOBAYASHI_EGFR_SIGNALING_24HR_DOWN), performed on gene array data obtained from PC9 cells treated with osimertinib (1 µM) or sorafenib (5 µM), alone or in combination (Combo), for two days. Enrichment scores (ES) and P values are reported. **B**, The data described in **A** were analyzed using our osimertinib-sorafenib combination signature (COMBO_DOWN). **C**, Heatmap and hierarchical clustering of the gene array data. **D**, Parental and gefitinib-resistant (EGFR-T790M, GR) PC9 cells were treated for 3 days with sorafenib (5 µM) or gefitinib (1 µM), alone or in combination, and immunoblot was performed using the indicated antibodies. **E**, PC9 cells containing highly complex CRISPR-barcodes were treated for two weeks with osimertinib (1 µM) and sorafenib (5 µM), alone or in combination (n=4 per condition). The percentage of barcodes enriched at least 5-fold in 1, 2, 3 or 4 replicates is shown.

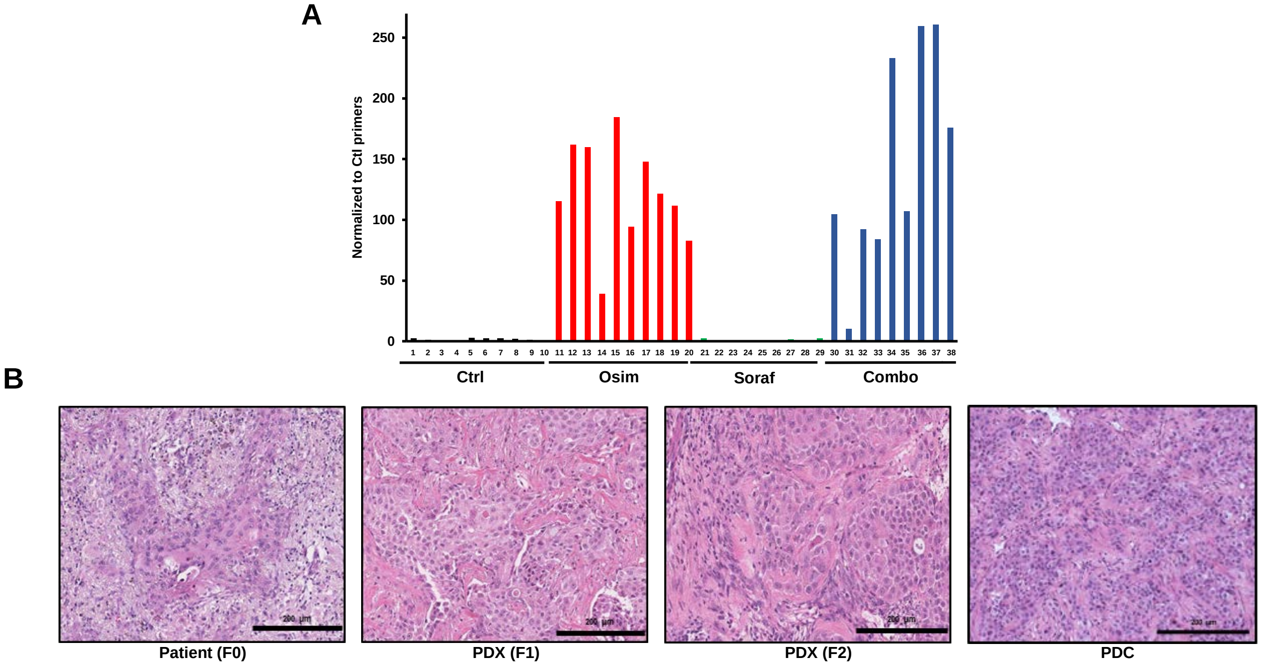


**Figure S7. A**, The levels of the EGFR-C797S CRISPR-barcode were measured by qPCR from genomic DNA derived from the tumors described in Fig. 6b. The values were normalized using control primers (the value of 1 corresponds to the levels measured in the injected cells). **B**, Representative images of hematoxylin-eosin staining of the YHIM-1024 original tumor, the corresponding YHIM-1024 PDXs (F1 inoculated in nude mice, F2 in NOG mice) and tumors obtained in SCID mice from YUX-1024 PDCs. Scale bar, 200 µm.

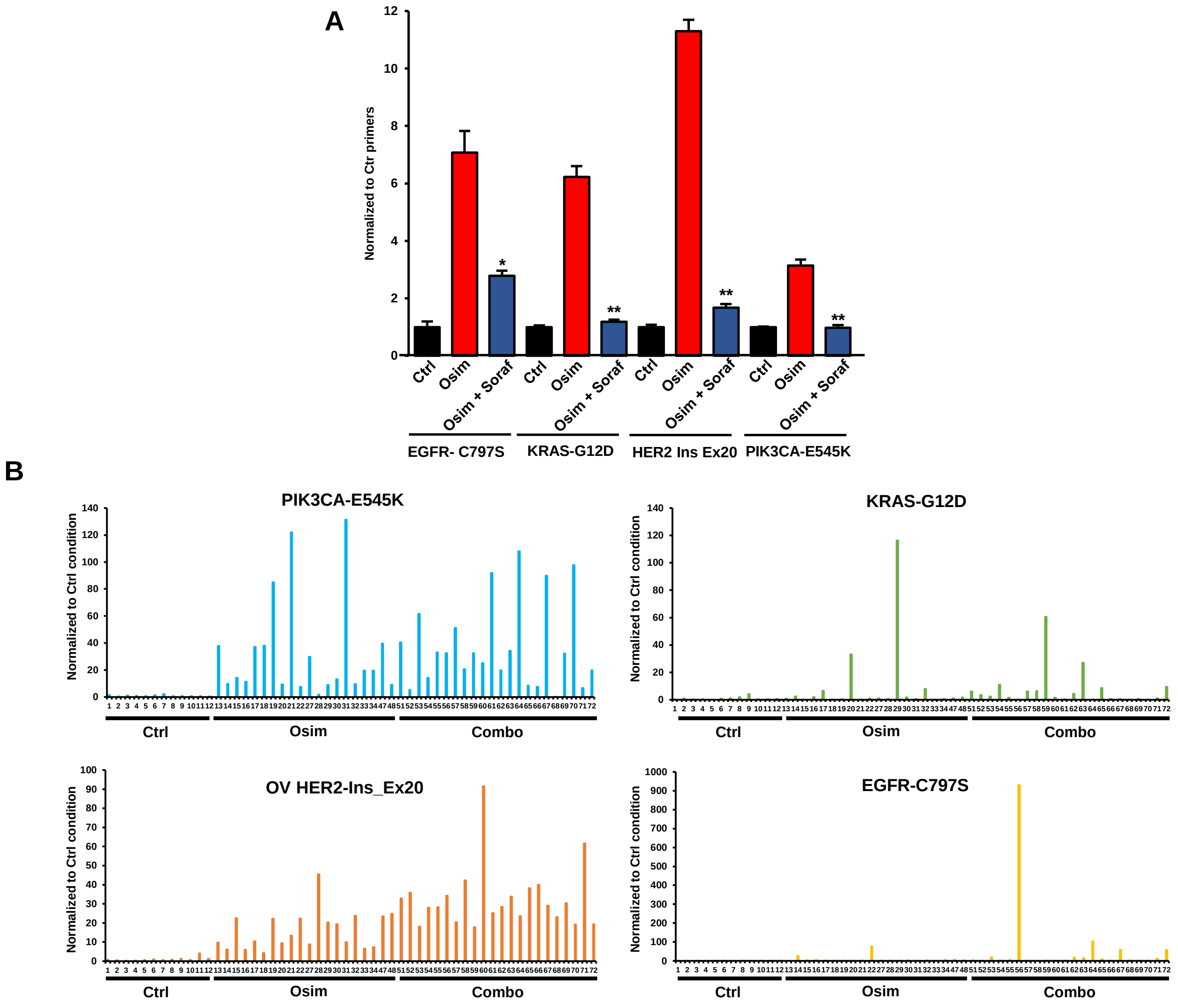


**Figure S8. A**, Effects of a 15-day treatment with osimertinib (0,1 µM) with or without sorafenib (5 µM) on the proportion of the indicated barcodes in the same multiplex PC9 cell model used for the mouse experiment described in Fig. 7a,b. The mean ±SEM (n=5) of one representative of three experiments is represented. B, The levels of the indicated barcodes were measured by qPCR from the tumors described in Fig. 7a,b.

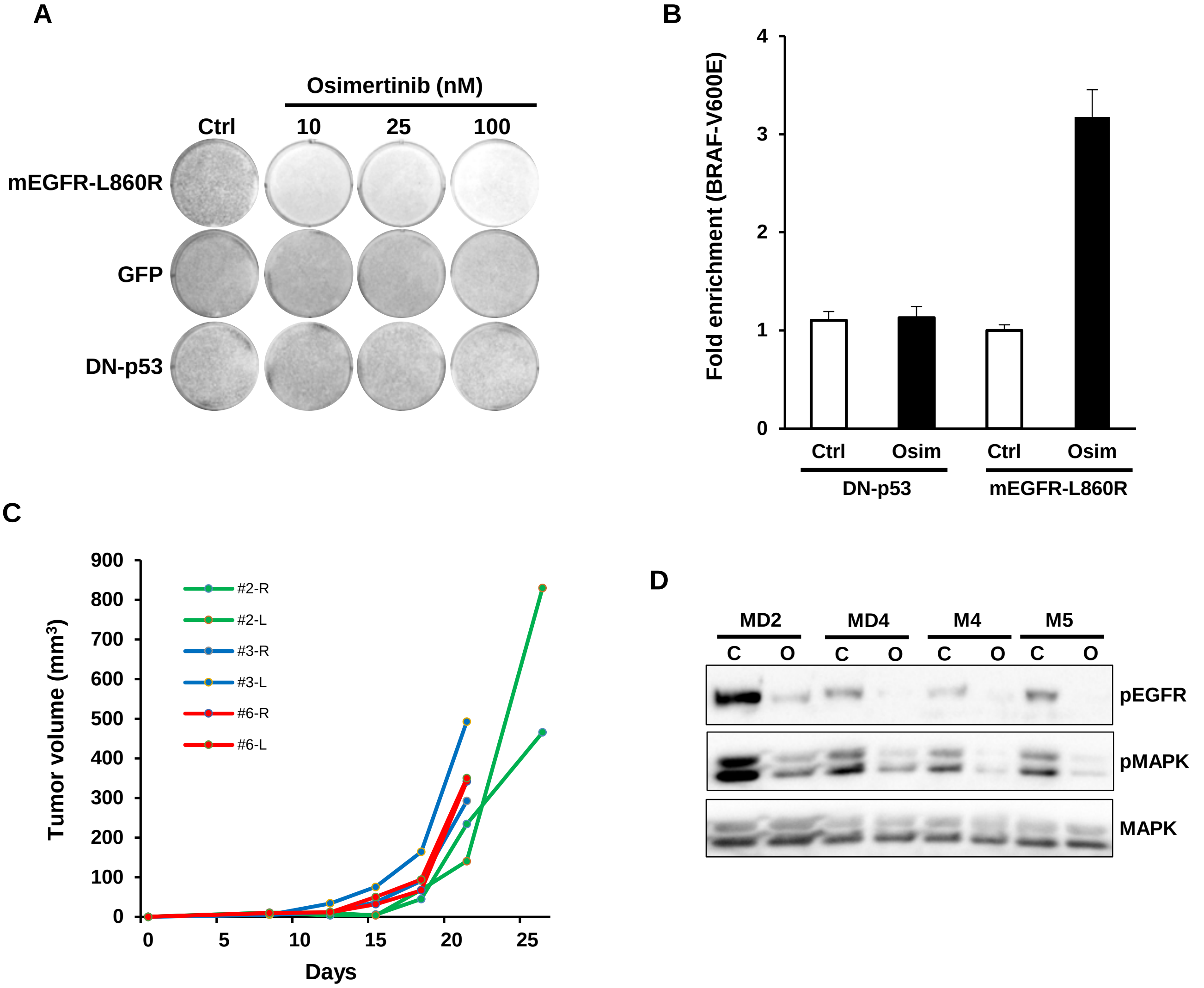


**E**

| | Control | Osim | Soraf | Combo |
|---|---|---|---|---|
| α-CD8 | 1 ± 0.3 | 16.5 ± 0.2 | 15.8 ± 2.9 | 16.9 ± 5.6 |
| α-CD3 | 12.4 ± 6 | 12.9 ± 7 | 13.3 ± 4.5 | 8.4 ± 4 |
| α-CD68 | 6.5 ± 1.9 | 23 ± 3.3 | 2.4 ± 0.4 | 34.5 ± 2.9 |
| α-CD86 | 14.2 ± 2.6 | 39.4 ± 11.7 | 26.7 ± 3 | 44.2 ± 6.9 |

**Figure S9. Development of a new syngeneic model of oncogenic addiction to mutant EGFR. A**, BALB-3T3 cells expressing mouse EGFR-L860R, GFP or DN-p53 were treated for a week in 1% calf serum with the indicated concentrations of osimertinib, then fixed and stained with crystal violet. **B**, A small subpopulation of BRAF-V600E cells was generated by CRISPR-barcoding in BALB-3T3-EGFR-L860R or DN-p53. The cells were grown for one week in the presence or the absence of osimertinib (100 nM) and the proportion of the BRAF-V600E barcode was measured by qPCR from genomic DNA. **C**, BALB-3T3-EGFR-L853R cells were subcutaneously injected in the right (R) and left (L) flanks of three BALB/c mice and the volume of the tumors was measured by caliper. **D**, One of the BALB-3T3-EGFR-L860R tumors shown in C was dissected and the cells grown in culture. Different individual clones were isolated (here 4 clones are shown) and treated for two hours in the presence (O) or the absence (C) of osimertinib 100 nM (1% calf serum), followed by western blot with the indicated antibodies. **E**, BALB/c mice bearing BEM-5 tumors were treated with osimertinib (20 mg/kg) and sorafenib (60 mg/kg), alone or in combination for ten days, followed by IHC using the indicated antibodies. The number of stained cells per field are indicated (mean ± SEM of four randomly chosen fields per tumor, three mice per condition).

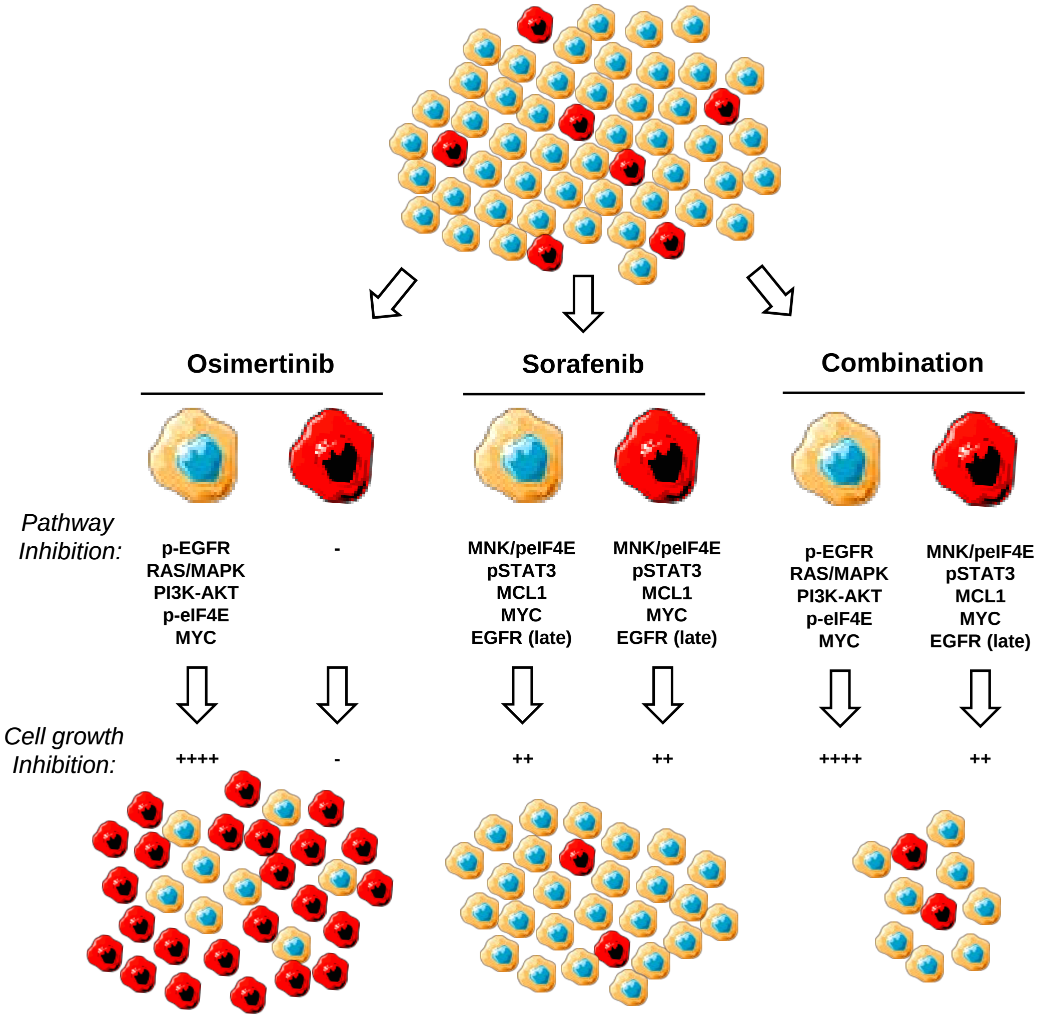


**Figure S10**. Diagram illustrating the effects of osimertinib and sorafenib, alone or in combination, on a mass population of osimertinib-sensitive (yellow) and osimertinib-resistant (red) NSCLC cells.
